# SUMO Inhibition Plus CD40 Agonism Increases Anti-Tumor Immunogenicity Through Interferon Mediated Macrophage Activation

**DOI:** 10.1101/2024.12.03.626688

**Authors:** Kevin Li, Asimina Courelli, Hyojae James Lee, Tatiana Hurtado de Mendoza, Alexei Martsinkovskiy, Evangeline Mose, Jay Patel, Izzy Ng, Siming Sun, Mohottige Don Neranjan Tharuka, Hervé Tiriac, Yuan Chen, Andrew M. Lowy

## Abstract

Resistance to immunotherapy is a cardinal feature of pancreatic ductal adenocarcinoma (PDAC). Inhibition of Small Ubiquitin-like MOdifier (SUMO), a post-translational modification with important immune regulatory functions, augments responsiveness to immunotherapy in non-PDAC models via pro-immunogenic effects on myeloid cells, cancer cells, and T-cells. Recently, it has been reported that SUMO inhibition has direct immunogenic effects on PDAC. Here, we report that the novel combination of SUMO inhibition with a small molecule, TAK-981, plus antibody-mediated CD40 agonism improves survival in an aggressive orthotopic mouse model of PDAC by enhancing anti-tumoral immunogenicity. This combination amplifies CD8+ T-cell tumor infiltration and induces significant changes among macrophages. TAK-981 also leads to enhanced cancer specific MHC-I expression both *in vitro* and *in vivo* by augmenting interferon signaling. We show that the improvement in survival is mediated by macrophages. Our findings show that SUMO inhibition complements CD40 agonism to enhance immune activity in PDAC via interferon signaling, improving survival in an aggressive pre-clinical model of PDAC and translating previous findings to a characteristically immunosuppressive and highly aggressive solid malignancy.

## Introduction

Pancreatic ductal adenocarcinoma (PDAC) is a lethal malignancy which confers a poor prognosis for most patients, despite recent improvement in outcomes with cytotoxic combination therapies.^1,2^ Immunotherapy has been largely ineffective, absent the small number of patients whose tumors harbor mismatch repair gene mutations. Though there are many reasons for PDAC’s poor immunogenicity, its mutational burden is modest, and it is widely recognized that the tumor immune microenvironment (TIME) of PDAC is characterized by fibrosis, infiltration of myeloid-derived suppressor cells (MDSC), regulatory T-cells, and tumor-associated macrophages (TAMs) that combine to inhibit infiltration and/or activity of CD8+ T-cells.^3^

The Small Ubiquitin-like MOdifier (SUMO) is a post-translational modification whose role in regulating innate and adaptive immunity has been greatly elucidated in recent years. SUMO knockout mice were previously shown to have heightened innate immune activity in response to pathogenic stimuli in a type I interferon-dependent manner.^4^ These effects extend beyond systemic adaptive immunity into the tumor immune microenvironment (TIME), where mice bearing either colorectal tumors or lymphoma, when treated with a small molecule inhibitor of SUMOylation, displayed profound increases in effector T-cell, dendritic cell, and anti-tumor macrophage activity, leading to successful combinations with immunotherapies.^5,6^ Importantly, in these studies, SUMO inhibition led to increased CD40 expression in human dendritic cells cultured *ex vivo*, as well as in mice bearing lymphomas. In a subcutaneous mouse model of PDAC, SUMO inhibition also leads to increased CD8+ T-cell tumor infiltration, suggesting that SUMO inhibition may also prime PDAC tumors for response to immunotherapy.^7^ This finding was recapitulated in an orthotopic mouse model of PDAC, where mice bearing orthotopically injected KPC tumors were treated with TAK-981, leading to increased T-cell tumor infiltration and T-cell dependent survival.^8^ In addition, single-cell RNA sequencing analysis of these tumors showed important pro-immunogenic changes in the tumor-associated macrophage population in a manner that is associated with IFNγ signaling.^8^ These studies also showed enhanced tumor necrosis factor (TNF) pathway activity in multiple cell types, including increased expression of TNF receptor family members OX-40 and 4-1BB in T-cells and CD40 in myeloid cells.^8^

In addition to its effects on tumor infiltrating leukocytes, SUMO inhibition has direct effects on cancer cells. Pharmacologic SUMO inhibition in hematologic cancer cells leads to increases in type I interferon signaling via stimulation of IFNβ secretion, an effect not previously seen in solid tumors.^5^ As a result, SUMO inhibition leads to increased STAT1 expression and the expression of downstream interferon-stimulated genes, including those necessary for antigen presentation via MHC-I.^9^ SUMO inhibition upregulates cytokine-dependent cancer specific MHC-I expression in DLBCL, osteosarcoma, neuroblastoma, breast cancer, and lung cancer lines,^9^ as well as in an orthotopic mouse model of PDAC.^8^ In multiple cancer models, downregulation of cancer specific MHC-I leads to decreased T-cell infiltration and immune escape,^10–13^ suggesting that cancer specific MHC-I is necessary for effective immunotherapy, and that inhibiting SUMO in cancer cells also leads to anti-tumor, immune-based effects. Though the effects of SUMO inhibition on the tumor immune microenvironment may be broad and largely pro-immunogenic, the benefit of combining SUMO inhibition with immunotherapy in PDAC has not yet been thoroughly explored.

Among the many immunotherapeutic agents under investigation for treating PDAC, anti-CD40 agonism is of particular interest.^14,15^ CD40 is a member of the TNF receptor family expressed on antigen presenting cells, such as dendritic cells, B cells, and macrophages, thereby playing a central role in pro-inflammatory cytokine secretion and T cell priming and activation.^16,17^ Combining conventional chemotherapy with αCD40 agonistic antibody led to radiographic responses in patients with metastatic PDAC via a T-cell independent mechanism.^15,18,19^ Though it increases T-cell tumor infiltration, αCD40 therapy functions primarily via its effects on myeloid and dendritic cells.^18,20^ Therefore, αCD40 agonistic antibody in combination with chemotherapy fundamentally alters the nature of the PDAC microenvironment, allowing for CD8+ T-cells to infiltrate tumors in a macrophage-dependent manner.

Because of the multi-faceted pro-immunogenic effects of SUMO inhibition established in non-PDAC and PDAC mouse models, as well as the myeloid-dependent effects of antibody-mediated CD40 agonism, we hypothesized that the novel combination of SUMO inhibition with the small molecule inhibitor TAK-981 plus αCD40 agonistic antibody would enhance antitumor immunogenicity in PDAC. Here, we first show that the combination of SUMO inhibition with CD40 agonism leads to improved overall survival in an aggressive orthotopic mouse model of PDAC. Next, using a combination of single cell RNA sequencing (scRNAseq) and flow cytometry, we show that this novel combination therapy increases antitumor immunogenicity within the TIME by increasing CD8+ T-cell infiltration, altering the macrophage population, and increasing cancer-specific MHC-I expression. We show that TAK-981 primes cancer cells to respond to interferon signaling *in vitro*, and that CD40 agonism leverages this priming effect on cancer cells *in vivo*. Finally, using a combination of antibody, genetic, and drug-based depletion strategies, we show that the survival advantage conferred by SUMO inhibition and CD40 agonism is dependent on macrophages rather than CD8+ T-cells.

## Results

### TAK-981 combined with CD40 agonism improves survival in PDAC bearing mice

Given that SUMO inhibition upregulates CD40 expression in myeloid cells,^8^ and that SUMO inhibition and CD40 agonism each cause activating effects on innate and adaptive immunity, we initially hypothesized that TAK-981 would enhance the efficacy of CD40 agonism to increase survival and decrease tumor burden in mice with PDAC. To test this hypothesis, we generated orthotopic PDAC tumors by injecting KPC46 cells into the pancreata of syngeneic mice. After confirming tumor growth via ultrasound, we treated mice with either TAK-981, αCD40 agonistic antibody alone, or in combination (**Fig 1A**). Combination therapy led to a significant increase in median survival compared to untreated mice (untreated median survival 19 days vs. TAK-981 + αCD40 median survival 24.5 days, p = 0.0005). However, compared to untreated mice, those treated with αCD40 alone did not have a median survival benefit (19 d vs. 20 d, p=0.417), nor did mice treated with TAK-981 alone (19 d vs. 15.5 d, p= 0.879) (**Fig 1B**). In an independent experiment, the tumors from mice sacrificed after 12-13 days of combination treatment weighed significantly less than those of vehicle-treated mice (vehicle + IgG: 1.10 g vs. TAK-981 + αCD40: 0.57 g, p= 0.01) (**Fig 1C**). Tumors in the combination treatment group were numerically smaller in diameter, though this difference did not reach statistical significance ( **Fig 1D**). Interestingly, despite these findings, there were no significant histologic changes on conventional hematoxylin and eosin staining associated with any of the treatments (**Fig 1E**). Taken together, these findings show that TAK-981 plus αCD40 combination therapy enhances median survival and decreases tumor burden in an aggressive orthotopic PDAC mouse model.

**Figure 1.**
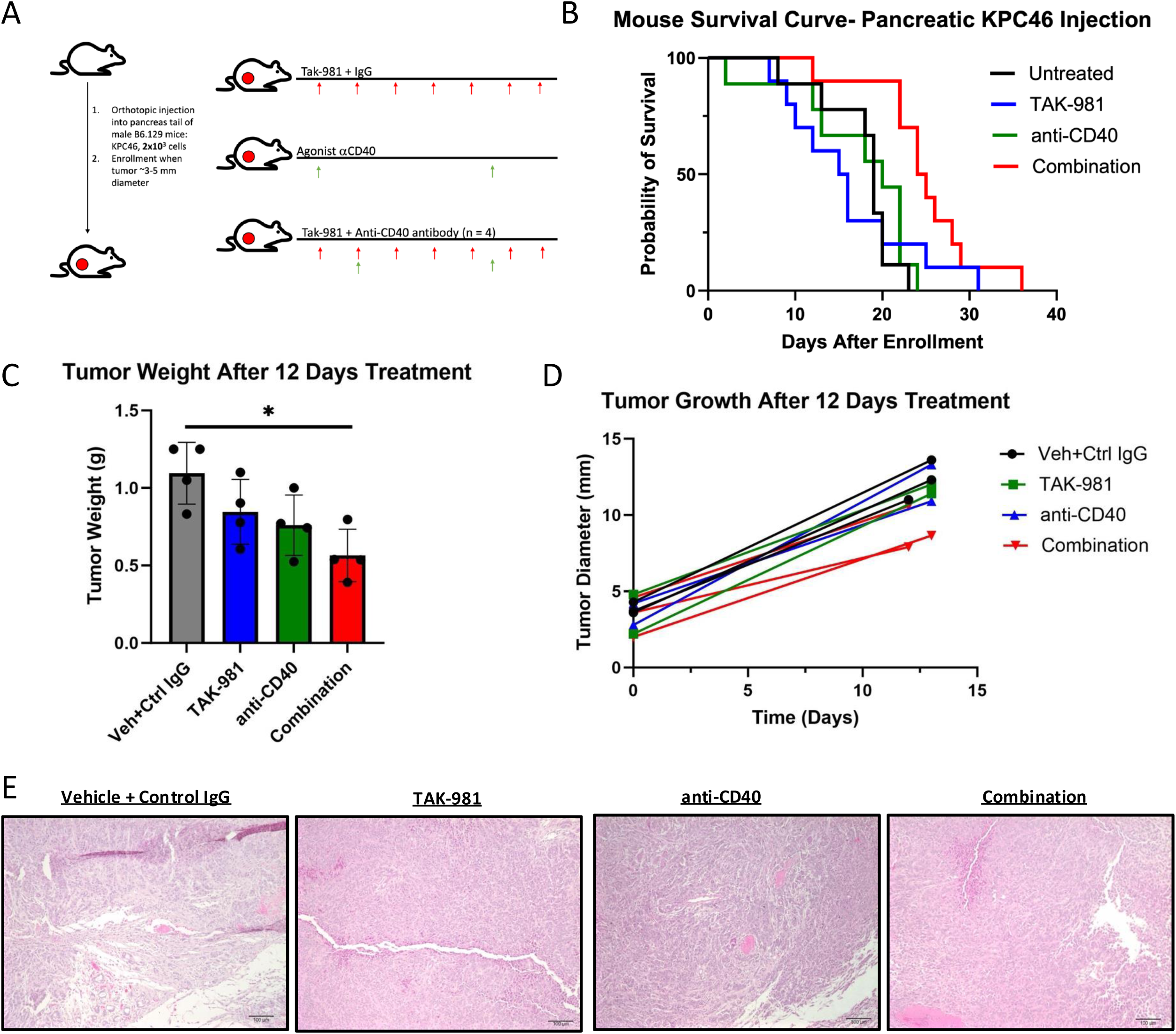
TAK-981 combined with CD40 agonism improves survival in an orthotopic mouse model of PDAC. A) Experimental timeline. Mice were injected with 2,000 KPC46 cells and enrolled into one of four groups: untreated, TAK-981, anti-CD40, and combination. Mice were enrolled and randomized when tumors reached 3-5mm in diameter as measured by ultrasonography. Mice were dosed with TAK-981 at 15mg/kg every 48 hours and αCD40 at 100µg every 7 days. B) Survival curves for untreated (median survival: 19 days), TAK-981(median survival: 15.5 days, p= 0.879), αCD40(median survival: 20 days, p=0.417), and combination (median survival: 24.5 days, p=0.0005). Survival curves were analyzed with the log rank test. C) Tumors were generated as in panel A and treated for 12 days with either vehicle + IgG control, TAK-981, αCD40, or TAK-981+αCD40. Tumor weights for vehicle + IgG (1.10 g), TAK-981(0.85 g, p=0.31), αCD40 (0.76 g, p=0.11), combination (0.57 g, p= 0.01). Weight comparisons to the vehicle + IgG group were conducted using a two tailed t-test. D) Mean tumor diameters for vehicle + IgG (12.3 mm), TAK-981 (11.13 mm, p=0.67), αCD40 (11.69, p=0.93), and combination (9.05mm, p=0.05). Comparisons to vehicle + IgG group were conducted using a two tailed t-test. E) Representative hematoxylin and eosin staining for tumors in the various treatment groups.

### Combination treatment induces broad pro-immunogenic changes in the tumor immune microenvironment

To gain insight into any changes in the tumor immune microenvironment induced by combination treatment, and to identify cell types primarily responsible for driving treatment efficacy, we analyzed tumors using single cell RNA sequencing (scRNAseq). We generated orthotopic tumors and treated mice with either TAK-981 alone, αCD40 alone, or in combination for 12-13 days after tumors reached 3-5mm in diameter as measured by ultrasound imaging. After this treatment period, we sacrificed the mice and processed the tumors for scRNAseq. We identified a broad variety of cell types, including cancer cells, T-cell subsets, myeloid cell subsets, neutrophils, NK cells, fibroblasts, and endothelial cells **(Fig 2A**). B-cells were not well-represented in this data set. The treatments induced distinct changes in the sizes of individual cell populations. Of note, combination treatment was associated with a large increase in the proportion of T-cells (Vehicle + IgG: 3.05%, TAK-981: 7.4%, αCD40: 1.4%, TAK-981 + αCD40: 10.8%; Effect Size for Vehicle + IgG vs TAK-981 + αCD40: 1.54) (**Fig 2B, 2C**). TAK-981 treatment alone caused a smaller, but measurable increase in tumor infiltrating T-cells (**Fig 2B, 2C**), recapitulating previously reported findings of T-cell infiltration into mouse PDAC tumors.^7,8^ In addition, combination treatment led to an increase in the proportion of macrophages compared to mice treated with monotherapy or vehicle controls (Vehicle + IgG: 7.1%, TAK-981: 8.9%, αCD40: 6.7%, TAK-981 + αCD40: 13.0%; Effect Size for TAK-981 + αCD40 vs. αCD40: 0.79) (**Fig 2B, 2D**).

**Figure 2.**
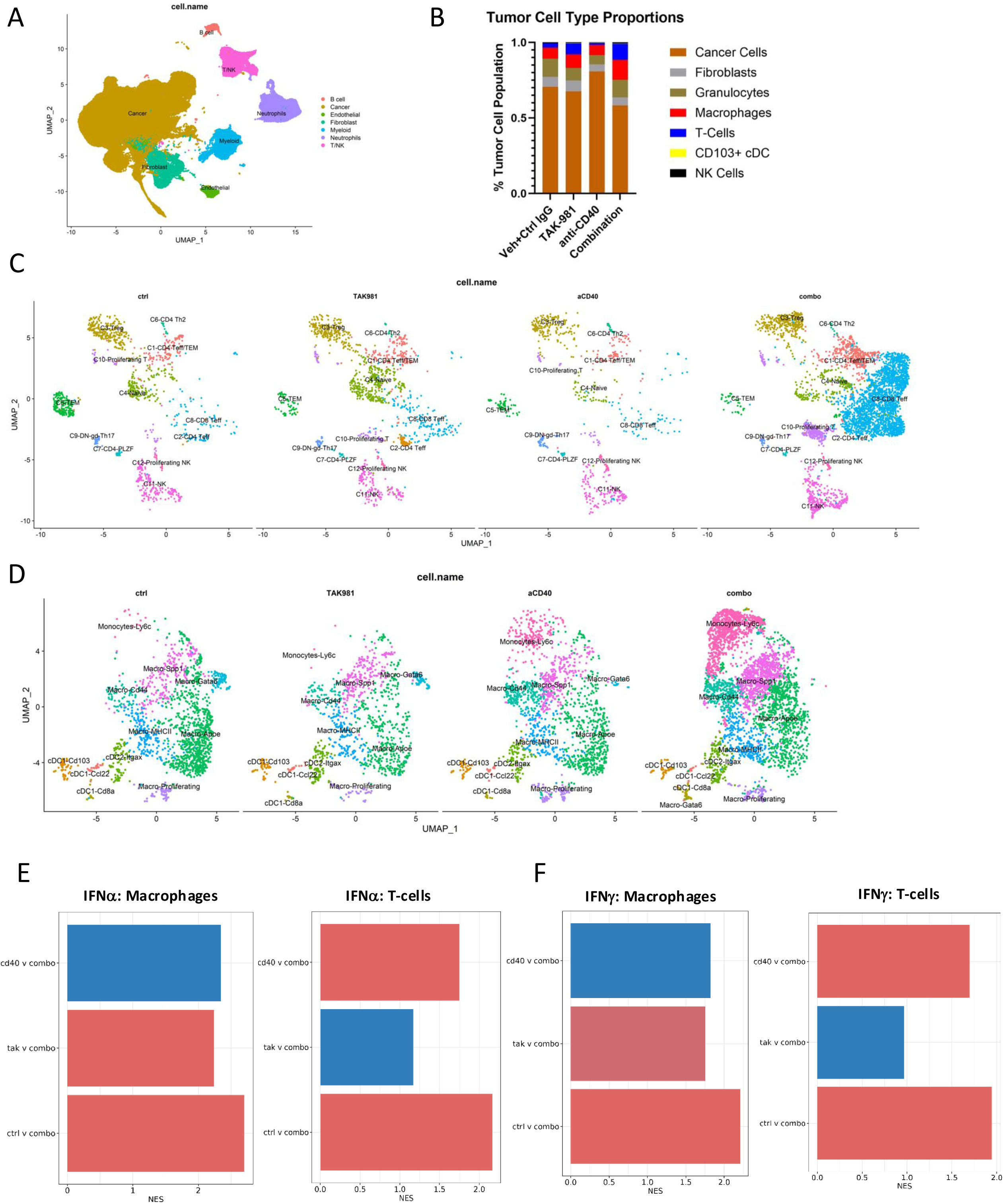
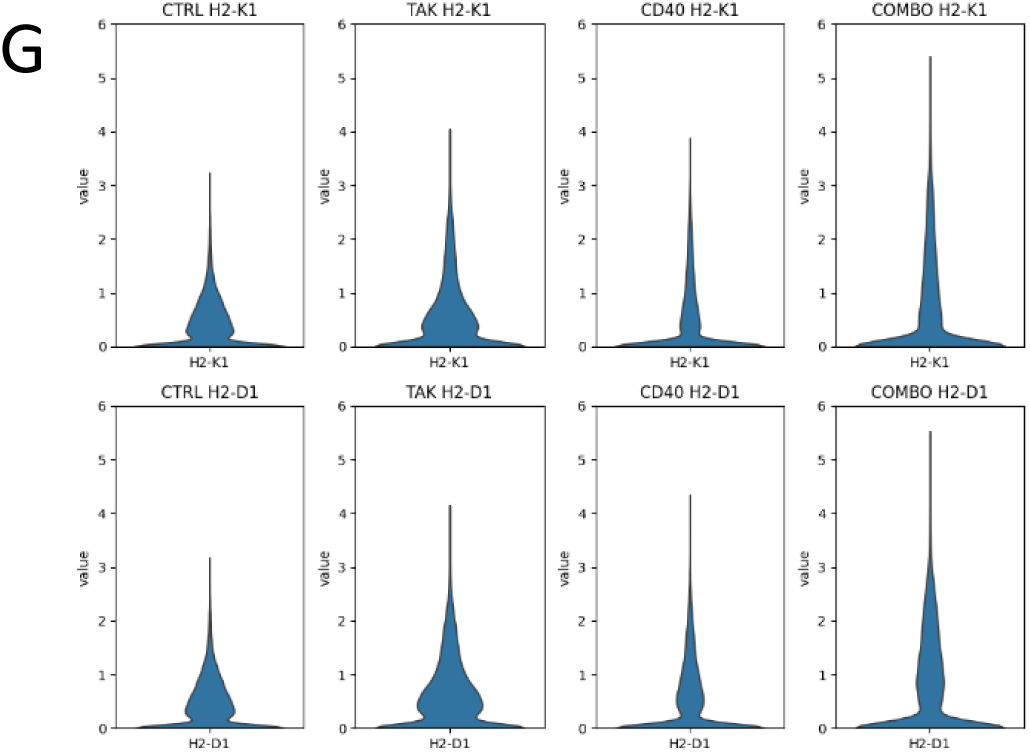
TAK-981 combined with CD40 agonism induces broad pro-immunogenic changes in the tumor immune microenvironment. A) Umap of tumor cell populations sorted by cell type (left) and treatment condition (right). B) Stacked bar graph of tumor cellular composition for each treatment condition. T Cell Quantification (Vehicle + IgG: 3.05%, TAK-981: 7.4%, αCD40: 1.4%, Combination: 10.8%, Effect Size for Combination vs Vehicle + IgG: 1.54 . Macrophage quantification (Vehicle + IgG: 7.1%, TAK-981: 8.9%, αCD40: 6.7%, combination: 13.0%, Effect Size for anti-CD40 vs Combination: 0.79). C) Umap of lymphoid immune cell populations per treatment condition. D) Umap of myeloid immune cell populations per treatment condition. E) GSEA Analysis Pairwise Comparisons for IFNα-Macrophages (Left): Combination vs αCD40 (NES: 2.33, p<0.0001), Combination vs TAK-981 (NES:2.24, p<0.0001), and Combination vs Vehicle + IgG (IFNα-NES: 2.70, p<0.0001). T Cells (Right): Combination vs αCD40 (NES: 1.75, p= 0.005), Combination vs TAK-981 (NES: 1.16, p=0.42), and Combination vs Vehicle + IgG (NES: 2.17, p<0.0001. F) GSEA Analysis Pairwise Comparisons for IFNγ-Macrophages (Left): Combination vs αCD40 (NES: 1.82, p=0.0002), Combination vs TAK-981 (NES: 1.75, p<0.0001), and Combination vs Vehicle + IgG (NES: 2.18, p<0.0001). T Cells (Right): Combination vs anti-CD40 (NES: 1.69, p=0.0009), Combination vs TAK-981 (NES: 0.97, p=0.99), and Combination vs Vehicle + IgG (-NES: 1.94, p <0.0001).G) Violin plots for MHC-I expression (H2-K1, H2-D1) for TAK+ αCD40, monotherapy, and vehicle + IgG conditions. Pairwise comparison differential expression coefficients for H2-K1 (Combination vs anti-CD40: 0.76, Combination vs TAK: 0.06, Combination vs Vehicle + Control IgG: 0.60; p<0.002 for all pairwise comparisons) and H2-D1 (Combination vs anti-CD40: 0.68, Combination vs TAK-981: 0.13, Combination vs Vehicle + IgG: 0.63; p<0.001 for all comparisons)

To gain insight into specific pathways that may contribute to these findings, we conducted gene set enrichment analysis (GSEA) on individual cell populations across treatment conditions. Based on prior reports that SUMO inhibition leads to upregulation of interferon signaling,^4,5^ we hypothesized that the additive benefit of combining SUMO inhibition with CD40 agonism may be due to increased immune cell infiltration via an additive increase in interferon signaling within the TIME, compared to monotherapy treatment. We conducted pairwise comparisons between treatment conditions for interferon α (IFNα) and interferon γ (IFNγ) signaling within T-cells and macrophages. In both T-cells and macrophages, IFNα- and IFNγ-mediated signaling were two of the highest differentially expressed signaling pathways. Pairwise comparisons between treatment conditions for interferon signaling in T cells showed significantly increased IFNα and IFNγ signaling in combination treatment versus αCD40 (IFNα: NES: 1.75, p= 0.005, IFNγ: NES: 1.69, p=0.0009) and combination versus control (IFNα: NES: 2.17, p<0.0001, IFNγ: NES: 1.94, p<0.0001). However, within T-cells, there was no change in IFNα and IFNγ signaling in combination versus TAK-981 alone (**Fig 2E, 2F**). Similar pairwise analysis for interferon signaling in macrophages showed a significant increase in IFNα and IFNγ increase in combination versus αCD40 (IFNα: NES: 2.33, p<0.0001, IFNγ: NES: 1.82, p=0.0002), combination versus TAK-981 alone (IFNα: NES: 2.24, p<0.0001, IFNγ: NES: 1.75, p<0.0001), and combination versus control (IFNα: NES: 2.70, p<0.0001, IFNγ: NES: 2.18, p<0.0001) (**Fig 2E, 2F**). We also found that combination treatment is associated with decreased TGFβ signaling between macrophages and fibroblasts, as well as between macrophages and epithelial cells (**Supplemental Figure S1**). Overall, these results show that TAK-981 alone drives an increase in IFNα and IFNγ in T-cells, consistent with prior reports in different *in vivo* cancer models,^5,7,9^ and that the addition of αCD40 leads to an additive increase in signaling through these pathways in both T-cells and macrophages.

Although combination treatment did not lead to similar global increases in IFNα and IFNγ signaling in cancer cells when compared to either monotherapy, they showed increased MHC-I expression when exposed to combination treatment *in vivo*. Since previous reports showed that TAK-981 increases cancer-specific MHC-I expression,^8,9^ we sought to understand whether changes in MHC-I expression in cancer cells occurred due to the treatment combinations. In our scRNAseq dataset, we quantified the expression of two key genes related to MHC-I expression: *H2K1* and *H2D1*. Expression of these genes modestly increased with TAK-981 alone and CD40 agonism alone. However, combination treatment led to a markedly larger increase in these genes (**Fig 2G**). Altogether, these findings show that SUMO inhibition by TAK-981 provides a priming effect on cancer cells, whereby it can potentiate interferon-dependent cancer cell specific MHC-I expression even in a highly aggressive, classically immunosuppressive tumor type.

### TAK-981 enhances type I interferon signaling and promotes cancer cell-specific MHC-I expression

Based on our scRNAseq analysis that showed an increase in cancer cell MHC-I expression in the tumors treated with combination therapy, we sought to recapitulate TAK-981’s priming effect on interferon signaling in cancer cells *in vitro.* To do this, we first explored the effects of TAK-981 in patient-derived organoid (PDO) models of PDAC. PDOs are a validated platform for assessing drug efficacy that faithfully recapitulate the tumor biology of the patient donors^21,22^ and provide a convenient platform to study biologically heterogeneous replicates. Though previous studies highlighted TAK-981’s cancer cell specific upregulation of type I interferon signaling in hematologic cancer cell lines, this was not previously observed in solid tumor cell lines.^5^ We first measured *in vitro* cytotoxicity as an indication of different PDO lines’ susceptibility to SUMOylation inhibitions, since interferon activation can lead to cell cytotoxicity in PDAC cancer cells.^23,24^ We found that TAK-981 induces cytotoxicity in a dose-dependent manner in several unique PDO lines, with an IC_50_ range of 16.5 nM to 116.9 nM (**Fig 3A**). GSEA^25^ of bulk RNA sequencing comparing untreated PDOs with TAK-981 treated PDOs showed significant increases in both IFNα and IFNγ responses after exposure to 100 nM TAK-981 for 48 hours (**Fig 3B**). Additionally, *in* vitro sensitivity to TAK-981 correlated with the degree of type I IFN response (**Supplemental Figure S2)**, suggesting that TAK-981 treatment not only leads to increased IFN signaling but also sensitizes cancer cells to extracellular IFN stimulation. HF44 and HF23 PDOs pre-treated with TAK-981 prior to exposure with IFNα generally showed an increase in downstream interferon stimulated genes compared to no pre-treatment with TAK-981 (**Fig 3C**). This effect was not seen in the HM1E PDO. This PDO also had the highest IC_50_, pointing to the differential sensitivity of cancer cells to TAK-981 mediated IFN upregulation. In conventional cell lines *in vitro*, TAK-981 treatment alone leads to a modest increase in STAT1 expression, as shown by RT-PCR quantification of STAT1 in KPC46 and 79E cells (**Fig 3D**). In all, these findings show that in a subset of PDAC cell lines, SUMO inhibition has a stimulatory effect on type I interferon signaling *in vitro*.

**Figure 3.**
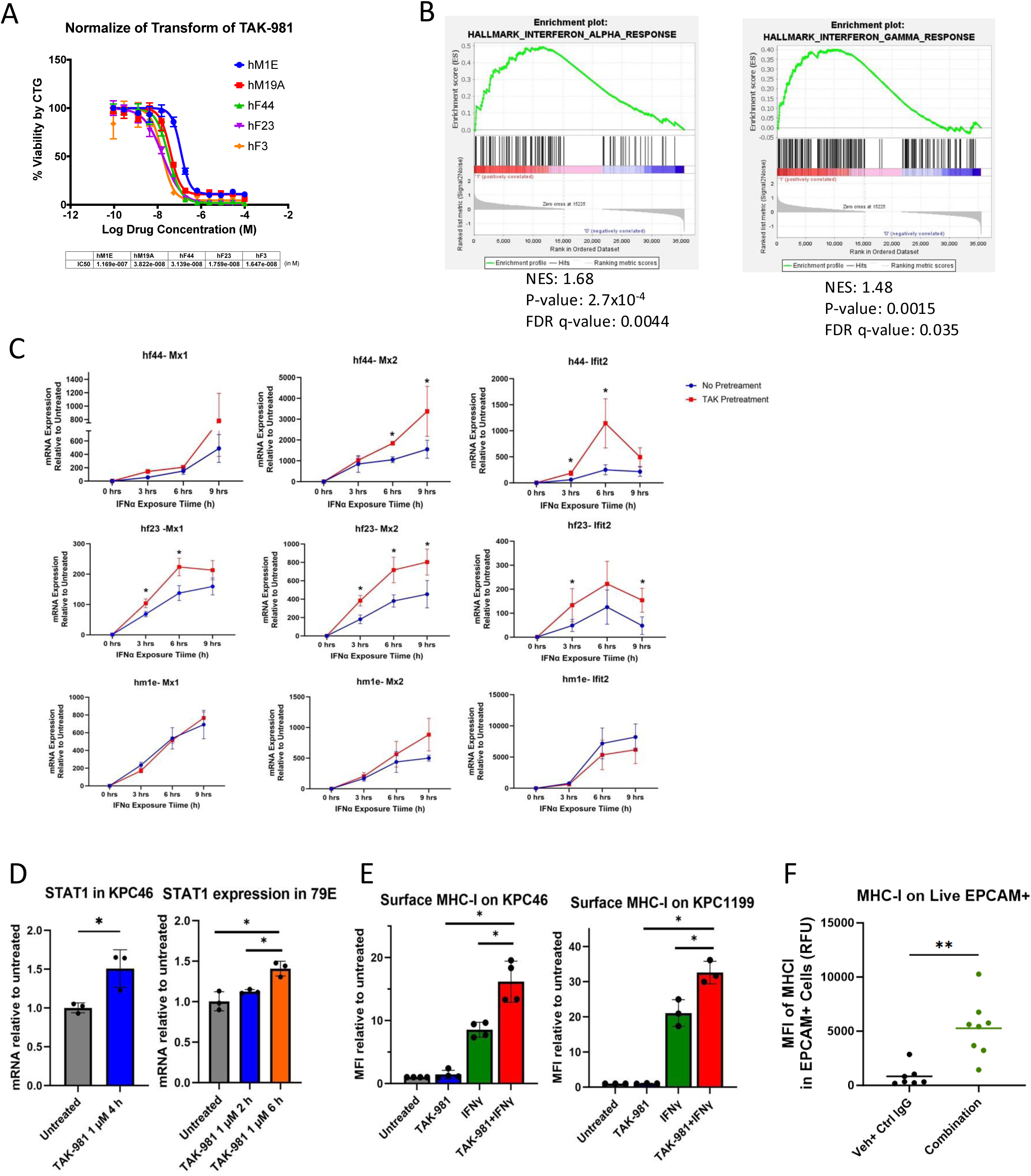
TAK-981 influences type I and type II interferon signaling to promote cancer cell-specific MHC-I expression. A) Dose response curves for five PDO lines, where cell viability was assessed after 5 days incubation with TAK-981 (dose range 16.5nM to 116.9 nM). B) Bulk RNA sequencing of untreated PDOs and PDOs exposed to 100 nM TAK-981 for 48 hours. Normalized enrichment plots showing increased IFNα (NES: 1.68, p<0.005) and IFNγ (NES: 1.48, p= 0.0015. C) Mx1, Mx2, Ifit2 PCR in hf44, hm1e, and hf23 PDO lines received either no pre-treatment or pre-treatment with 100 nM TAK-981 for 48 h followed by 1,000 U/mL IFNα (* p<0.05). D) STAT1 PCR in KPC46 and 79E following TAK-981 1µM exposure for either 2h, 4h or 6h. Data were normalized against the untreated condition. Statistical comparisons were conducted using a two tailed t test. KPC 46 STAT1: untreated: 1.0, TAK-981 1µM 4h: 1.51, p=0.03. 79E STAT1: untreated: 1.0, TAK-981 1µM 2h: 1.12 (p=0.02), TAK-981 1µM 6h: 1.41 (p=0.003). Individual points represent experimental replicates. E) Flow cytometry for MHCI in KPC46 and KPC 1199 following *in vitro* TAK-981 and IFNγ exposure mean fluorescent index (MFI) normalized to untreated condition: KPC 46 (untreated:1.00, TAK-981:1.42, IFNγ: 8.53, TAK-981+IFNγ:16.15, p<0.001 for pairwise comparisons with TAK-981+IFNγ). KPC 1199 (untreated:1.00, TAK-981:1.03, IFNγ:21.07, TAK-981+IFNγ:32.58, p<0.001 for pairwise comparisons with TAK-981+IFNγ). Replicates are individual experiments each normalized to its own negative control. F) Flow cytometry for MHCI on EPCAM+ tumor cells from Vehicle + Control IgG and Combination treated mice. Vehicle+Control IgG: 826 RFUs, Combination: 5269 RFUs, p = 0.001.

We sought to determine if a similar mechanism of increased IFN sensitivity following TAK-981 treatment could also explain the increased MHC-I expression we previously observed in scRNAseq analysis of *in vivo* PDAC tumors. Using flow cytometry, we measured surface level MHC-I expression on KPC46 cells cultured *in vitro* and overall found substantial additive increases in MHC-I expression when cells were exposed to both TAK-981 and IFNγ *in vitro* (MFI increase relative to untreated cells: TAK-981: 1.42-fold, IFNγ: 8.53-fold, TAK-981 + IFNγ: 16.15-fold, p<0.001 for pairwise comparisons with TAK-981 + IFNγ). Similarly, MHC-I expression on a separate mouse PDAC cell line, KPC1199, showed similar increases upon co-treatment with both TAK-981 and IFNγ (MFI relative to untreated: TAK-981: 1.03-fold, IFNγ: 21.07-fold, TAK-981 + IFNγ: 32.58-fold, p<0.001 for pairwise comparisons with TAK-981+IFNγ) (**Fig 3E**). These results show that TAK-981 can function to increase key machinery necessary for antigen presentation by PDAC cells by potentiating an IFNγ response. These findings were recapitulated *in vivo,* where *w*e processed tumors treated with either vehicle or TAK-981 + αCD40 combination therapy and selectively measured MHC-I expression on EPCAM+ cells. Tumors treated with combination therapy had EPCAM+ cells with significantly higher MHC-I expression compared to EPCAM+ cells in untreated tumors (Vehicle + IgG: 826 RFUs, TAK-981 + αCD40: 5269 RFUs, p = 0.001) (**Fig 3F**). Taken altogether, our findings show that TAK-981 enhances PDAC cells’ ability to respond to type I and II IFN, leading to synergistic increases in downstream gene expression and antigen presentation *in vitro*, and that the *in vivo* addition of CD40 agonism leverages this effect.

### TAK-981 plus αCD40 combination therapy increases CD8+ T-cell tumor infiltration

Since we observed increased T cell infiltration and MHC-I expression on cancer cells, we sought to explore whether combination therapy has detectable effects on CD8+ T-cells. scRNA sequencing analysis showed a diverse collection of T-cell subpopulations, among which there was a robust increase in the percentage of CD8+ T-cells only in tumors treated with combination therapy (**Fig 2C**). This increase in CD8+ T-cells tumor infiltration was verified with immunofluorescent staining of untreated and treated tumors taken at time of euthanasia (**Fig 4A and 4B**). Flow cytometric analysis of tumors treated with combination therapy for 12-13 days also demonstrated an overall increase in CD3+ T-cell infiltration in tumors treated with combination therapy (**Fig 4C**). Moreover, there was a significantly higher proportion of CD8+ cells among the total CD3+ population in tumors treated with combination therapy compared to those treated with monotherapy and no treatment (vehicle + IgG: 1.6%, TAK-981: 0.7%, anti-CD40: 2.17%, Combination: 10.74%, p<0.01 for all pairwise comparisons to Combination group) (**Fig 4C**). No changes in the proportion of CD4+ T-cells among all CD3+ cells were detected (**Fig 4D**), suggesting that TAK-981 plus αCD40 therapy preferentially enhances recruitment of CD8+ T-cells. Simultaneously, there was a decrease in CD25+/FoxP3+ regulatory T-cells (vehicle + IgG: 45.05%, TAK-981: 52.33%, anti-CD40: 43.70%, Combination: 18.40%, p<0.001 for Combination vs Vehicle + IgG and TAK-981) (**Fig 4D**). This may be the result of less CD4+ T-cell to regulatory T-cell differentiation, since the percentage of total CD4+ T-cells was not significantly affected. These results show that combination treatment induces changes in tumor-specific adaptive immunity by causing increased infiltration and activation of CD8+ T-cells while simultaneously decreasing the presence of immunosuppressive regulatory T-cells.

**Figure 4.**
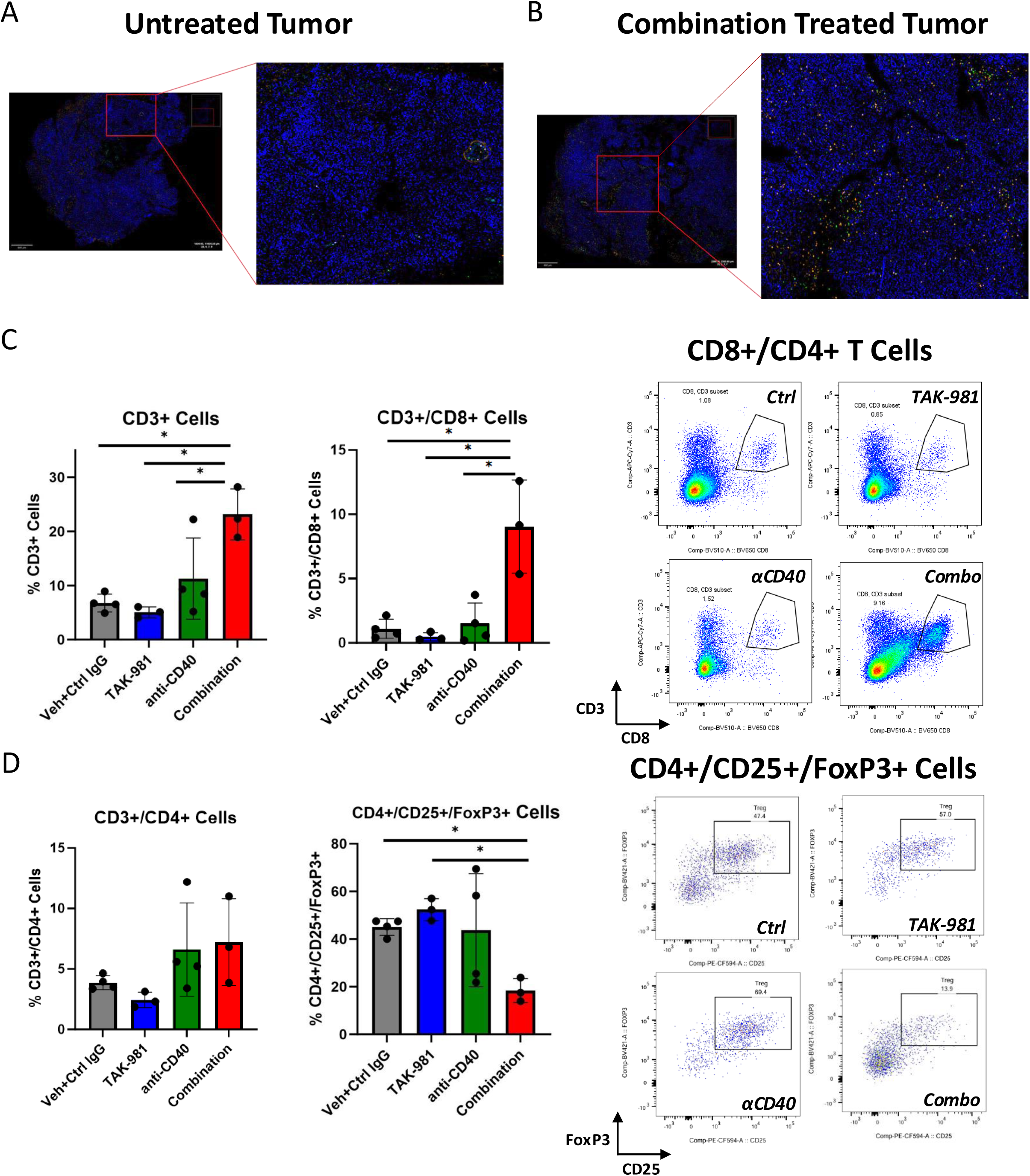
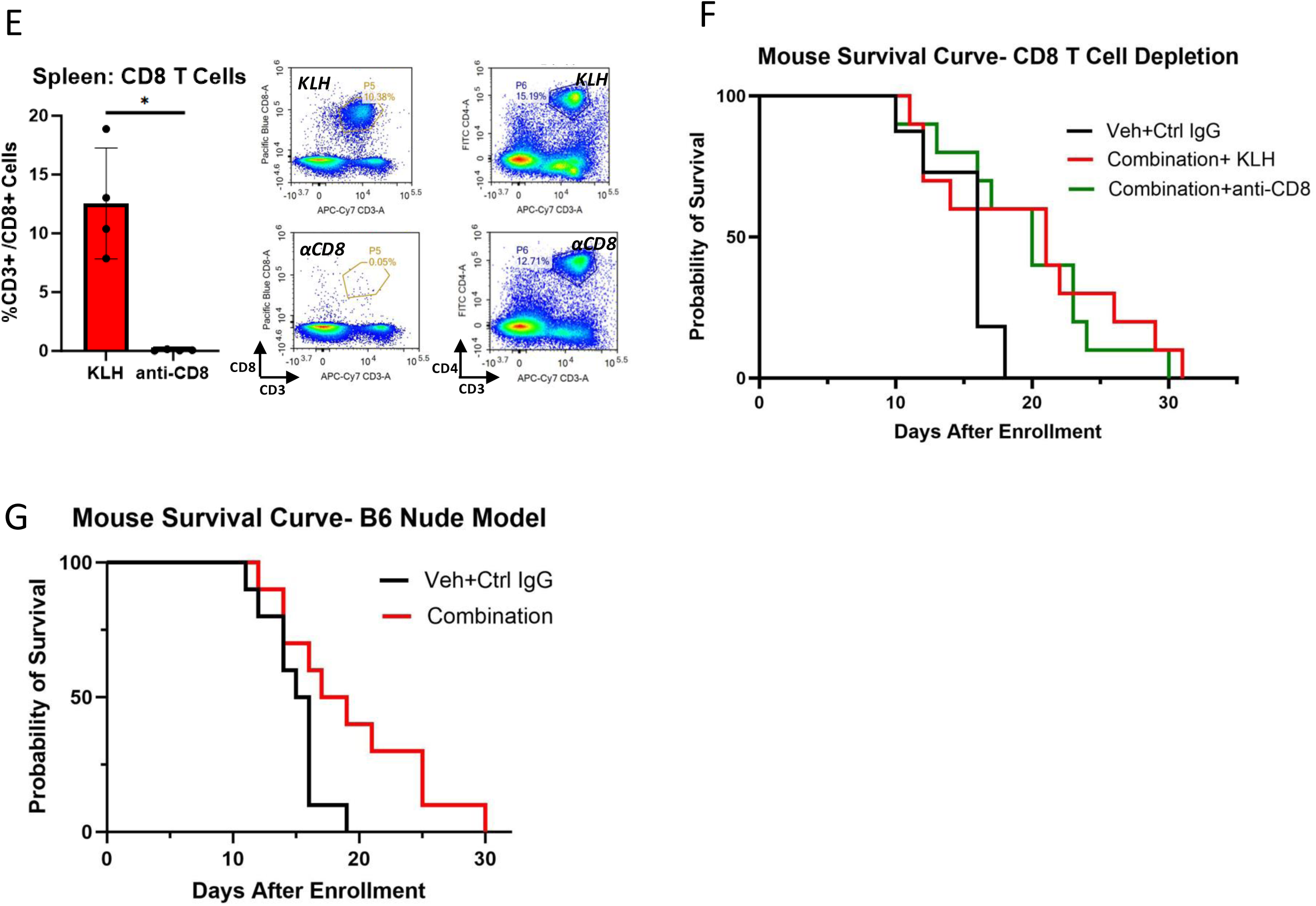
TAK-981 plus aCD40 agonism induced CD8+ T-cell infiltration does not mediate survival advantage: A) Representative immunofluorescence images of CD8 (orange) and CD4 (green) T cell infiltration in an untreated orthotopically generated tumor taken at the time of the animal’s euthanasia. B) Representative immunofluorescence images of CD8 (orange) and CD4 (green) T cell infiltration in an orthotopically generated tumor exposed to combination treatment and taken at the time of the animal’s euthanasia. C) T cell flow cytometry of tumors treated with vehicle + IgG, TAK-981, αCD40, or TAK-981 + αCD40 after 12 days of treatment. (Left) Flow cytometry quantification for CD3+ T Cells (vehicle + IgG:6.77%, TAK-981: 5.05%, αCD40: 11.28%, combination: 23.13%, p<0.02 for all pairwise comparisons to combination group), CD8+ T Cells (vehicle + IgG:1.08%, TAK-981: 0.48%, αCD40: 1.50%, combination: 9.04%, p<0.01 for all pairwise comparisons to combination group). (Right) Representative flow cytometry plots for each condition. D) (Left) Flow cytometry quantification for CD4+ T Cells (vehicle + IgG: 3.87%, TAK-981: 2.43%, αCD40: 6.61%, combination: 7.22%, ANOVA p= 0.14), TReg Cells (vehicle + IgG: 45.05%, TAK-981:52.33%, αCD40: 43.70%, combination: 18.40%, p<0.001 for combination vs vehicle + IgG and TAK-981. (Right) Representative flow cytometry plots for each condition. E) Spleen flow cytometry after 12 days of CD8 depleting antibody with sample flow cytometry plots for CD8 and CD4 T Cells (CD8 T Cells: KLH: 12.54%, αCD8: 0.05%, p<0.001). F) Survival curves for Selective CD8 T Cell Depletion. Tumors were generated via orthotopic injection of 2,000 KPC46 cells into mouse pancreases, and treatments were started after tumors reached 3-5 mm in diameter as measured via ultrasound imaging. Vehicle + IgG (median survival: 16 days), Combination + KLH isotype control (median survival: 21 days), Combination + αCD8 depletion antibody (median survival: 20 days). Survival comparisons were done with Log Rank test. Vehicle + IgG vs TAK-981 + αCD40 + CD8 depletion p = 0.04. Combination + KLH vs TAK+ αCD40 +CD8 depletion p = 0.67. G) Survival curves for B6 Nude Mice bearing orthotopically injected KPC46 pancreatic tumors either treated with vehicle + IgG controls (median survival 14 days) or TAK-981 + αCD40 combination (median survival 17 days), p= 0.032 as computed by the log rank test.

We then questioned whether the quantitative influx of CD8+ T-cells and decrease of regulatory T-cells, was responsible for driving the TIME toward a pro-immunogenic state and hypothesized that TAK-981 plus αCD40 combination therapy was functioning via a T-cell dependent mechanism. To test this hypothesis, we treated tumor bearing mice with TAK-981 plus αCD40 and used αCD8 antibody mediated depletion to eliminate CD8+ T-cells (**Fig 4E**).^26,27^ Mice treated with TAK-981 plus αCD40 and depleted of CD8+ T-cells did not experience significantly shorter survival compared to those receiving combination treatment but not CD8+ T-cell depletion (Vehicle + IgG median survival: 16 days, Combination + αKLH median survival: 21 days, Combination + αCD8 median survival: 20 days, Combination + KLH vs Combination + αCD8 depletion p = 0.67) (**Fig 4F**). In B6 nude mice devoid of an adaptive immune system, TAK-981 plus αCD40 conferred a modest, but statistically significant survival benefit (vehicle + IgG median survival: 14 days, Combination median survival: 17 days, p = 0.032) (**Fig 4G**). These results overall demonstrate that the therapeutic effects of TAK-981 plus αCD40 combination therapy are CD8+ T-cell independent. Taken altogether, while TAK-981 plus αCD40 combination therapy increases CD8+ T-cell tumor infiltration, these cells may not harbor sufficient functional activity to drive anti-tumor immunity.

### Clodronate-Mediated Macrophage Depletion Eliminates Survival Advantage and Tumor-Specific CD8+ T-cell Activation Conferred by Combination SUMO inhibition and CD40 Agonism

To explore the contribution of myeloid cells to the survival benefit of combination TAK-981 plus αCD40, we first verified that tumors treated with TAK-981 and αCD40 antibody had increased macrophage infiltration. Similar to our scRNASeq findings, which showed increased macrophage infiltration in mice treated with combination therapy, F4/80 staining in untreated and combination treated tumors from B6 nude mice also showed an increase in macrophage infiltration (**Fig 5A**). We then systemically depleted myeloid cells with clodronate in mice bearing orthotopic PDAC tumors.^18^ Myeloid depletion eliminated the survival advantage conferred by TAK-981 and αCD40 antibody (vehicle + IgG median survival: 13.5 d vs. Combination + PBS liposome median survival: 23 d, p = 0.01; vehicle + IgG: 13.5 d vs. Combination + Clodronate: 14 days p = 0.39) (**Fig 5B**). Flow cytometry analysis of spleens taken from the same tumor-bearing mice showed that clodronate was effective in eliminating macrophages and dendritic cells (**Fig 5C**). In addition, flow cytometry analysis of tumors showed that clodronate inhibited treatment-induced tumor infiltration of macrophages (vehicle + IgG: 11.9%, Combination + PBS liposome: 22.5%, Combination + Clodronate: 12.6%, p = 0.014) and dendritic cells (vehicle + Control IgG: 63.2%, Combination + PBS liposome: 55.6%, Combination + Clodronate: 14.9%, p<0.01) (**Fig 5D**). Interestingly, myeloid cell inhibition did not affect CD8+ T-cell infiltration. However, it did decrease CD8+ T-cell activation without affecting PD1 expression (**Fig 5E**). This suggests an interplay between myeloid cells and CD8+ T-cells whereby myeloid cells do not affect CD8+ T-cell tumor infiltration, but they play a necessary role in tumor-specific CD8+ T-cell activation in tumors treated with SUMO inhibition and CD40 agonism. Previous studies showed that PDAC bearing mice treated with chemotherapy and agonistic αCD40 antibody had decreased fibrosis and collagen deposition that were in turn driven by changes in macrophages.^20^ In our orthotopic model treated with SUMO inhibition and CD40 agonism, there were no changes in collagen deposition (**Supplemental Figure S3**). In all, these results show that macrophages are necessary for the survival and tumor-specific immunogenic benefits caused by the combination of SUMO inhibition and CD40 agonism.

**Figure 5.**
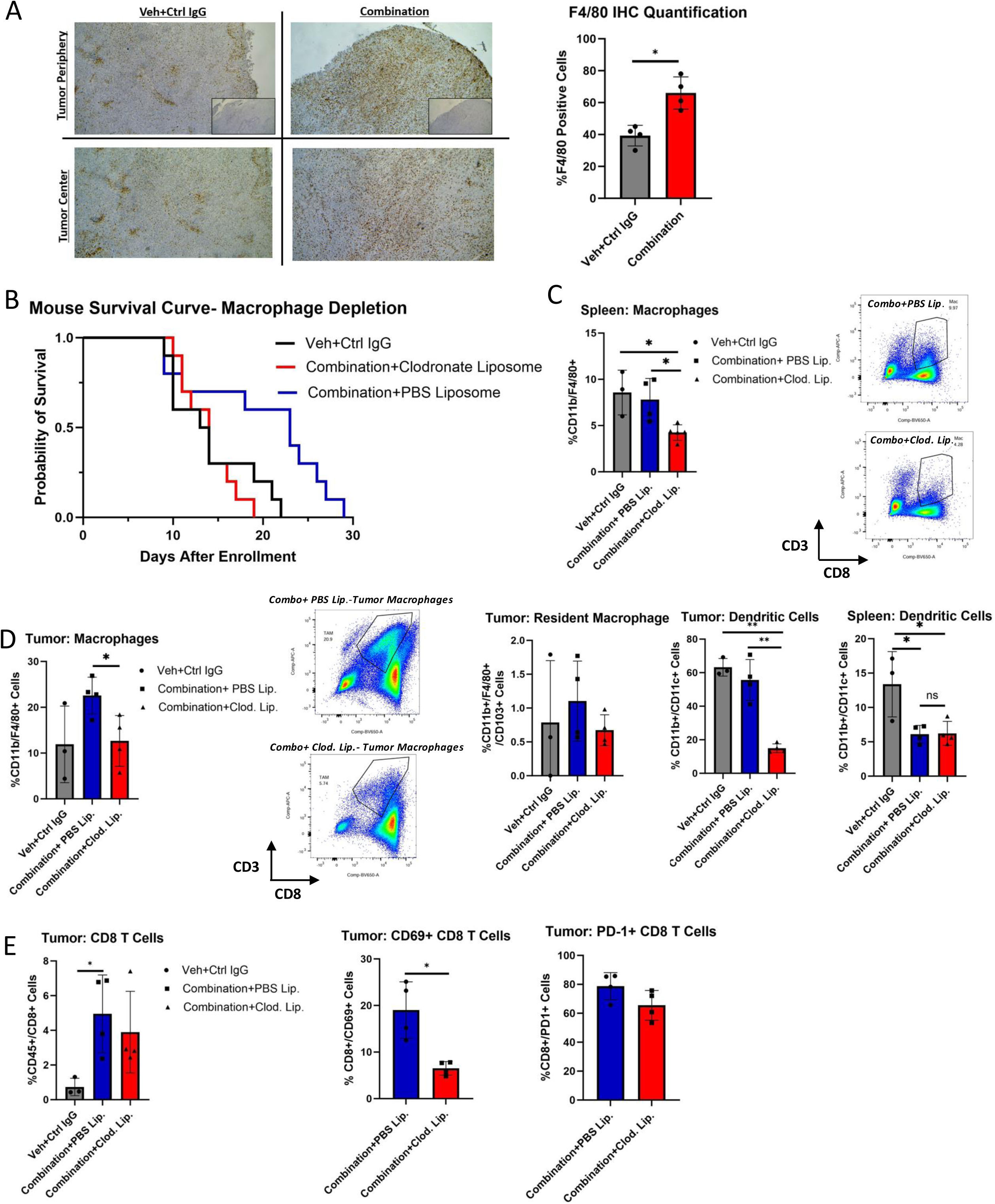
The increase in median overall survival caused by SUMO inhibition and CD40 agonism is dependent on myeloid cells: A) Immunohistochemistry staining for F4/80 and corresponding quantification in B6 nude mouse tumors treated with vehicle + IgG (39.5%) or TAK-981 +αCD40 (65.95%), p=0.008. B) Survival curves for clodronate-mediated macrophage Depletion. Vehicle + IgG (median survival: 13.5 days), TAK-981 + αCD40 +PBS liposome (median survival: 23 days), TAK-981 + αCD40 + clodronate liposome (median survival: 14 days). Survival comparisons with Log Rank test: Vehicle + IgG vs TAK-981 + αCD40 + PBS liposome, p = 0.01; Vehicle + IgG vs. TAK-981 + αCD40 + clodonrate liposome p = 0.39. C) (Left) Flow cytometry for spleen macrophages (vehicle + IgG: 8.57 %, TAK-981 + αCD40 + PBS liposome: 7.8 %, TAK-981 + αCD40 + clodronate liposome: 4.24%, p=0.01) with p-values comparing PBS liposome to clodronate liposome group. (Right) Representative flow cytometry plots for PBS liposome and clodronate liposome groups. D) Flow cytometry for tumor-associated macrophages with representative flow cytometry plots (vehicle + IgG: 11.9%, TAK-981 + αCD40 + PBS: 22.5%, TAK-981 + αCD40 +Clodronate: 12.6%, p=0.014), tumor resident macrophages (vehicle + IgG: 0.78%, TAK-981 + αCD40 +PBS: 1.10%, TAK-981 + αCD40 + clodronate: 0.67%, p=0.11), tumor dendritic cells (vehicle + IgG: 63.2%, TAK-981 + αCD40 + PBS.: 55.6%, TAK-981 + αCD40 + clodronate: 14.9%, p<0.01), and splenic dendritic cells (vehicle + IgG: 13.4%, TAK-981 + αCD40 +PBS: 6.2%, TAK-981 + αCD40 + clodronate: 6.2%, p=0.2) with p-values comparing PBS liposome to clodronate liposome groups. E) Flow cytometry for tumor specific CD8+ T cells (vehicle + Control IgG: 1.07%, TAK-981 + αCD40 + PBS: 4.95%, TAK-981 + αCD40 + clodronate: 3.85%, p=0.25), CD69+ CD8+ T Cells (TAK-981 + αCD40 + PBS: 39.5%, TAK-981 + αCD40 + clodronate: 15.6%, p=0.01), PD-1+ CD8+ T cells (TAK-981 + αCD40 + PBS: 78.7%, TAK-981 + αCD40 + clodronate: 65.5%, p=0.052) with p-values comparing PBS liposome to clodronate liposome groups.

## Discussion

Pancreatic cancer is marked by an immunosuppressive microenvironment that is characterized by infiltration of MDSCs and regulatory T-cells, as well as exclusion of cytotoxic T-cells.^3,28^ Within the PDAC TIME, myeloid cells may function to suppress immune reactivity by T-cell suppression and to upregulate inhibitory checkpoint proteins.^3^ As a result, PDAC is largely resistant to immunotherapy, with less than 5% of patients responding to anti-PD-1 monoclonal antibody therapy.^29,30^ Here we present a novel strategy of combining SUMO inhibition with CD40 agonism which increased overall survival in a highly aggressive orthotopic, syngeneic PDAC model. We found that SUMO inhibition and CD40 agonism lead to measurable pro-immunogenic changes in tumor-specific T-cell and macrophage populations. SUMO inhibition primes cancer cells *in vitro* and PDAC tumors *in vivo* to upregulate the strength of type I and type II interferon signaling, leading to increased MHC-I expression. Combination treatment also leads to concomitant increases in immunogenic macrophage and CD8+T-cells *in vivo*. Using both antibody-mediated and genetic means of T-cell depletion, we showed that treatment efficacy is preserved in mice depleted of CD8+ T-cells. However, treatment efficacy was abolished in mice exposed to the macrophage-depleting agent clodronate. As a result, this survival benefit is induced by a macrophage, not T-cell, dependent, mechanism.

Novel strategies to increase T-cell activation in PDAC have been studied preclinically, and in some instances these strategies successfully combined with immune checkpoint blockade. For instance, inhibiting autophagy in an orthotopic PDAC mouse model induced upregulation of MHC-I expression, which in turn increased T-cell activation and tumor infiltration. This effect sensitized the tumors to ICB.^13^ In a separate study, STAT3 inhibition increased T-cell activity and improved ICB efficacy.^26^ SUMOylation was previously shown to be an important regulator of adaptive immunity,^4,5^ and that SUMO inhibition can lead to potent immunogenic T-cell responses in mouse models of colorectal cancer and lymphoma.^5^ SUMO inhibition also increases cancer-specific MHC-I expression, an effect which is associated with ICB efficacy.^9^ Furthermore, SUMO inhibition decreased tumor growth in a CD8+ T-cell dependent manner in a subcutaneous tumor model of PDAC^7^ as well as an orthotopic PDAC mouse model.^8^ The results of these studies led us to initially hypothesize that rationally combining TAK-981 with specific immunotherapeutic agents might enhance anti-tumor T-cell activity, leading to improvements in overall survival in mouse models. We decided to investigate CD40 agonism because of its previously described immunogenic effects on T-cell activity in mouse models of PDAC.^18^ We reasoned that CD40 agonism may have an additive effect with SUMO inhibition because the latter has been previously shown to also exert direct effects on macrophages in an interferon-dependent mechanism.^6^

Interestingly, combining SUMO inhibition with CD40 agonism led to pro-immunogenic effects in both adaptive and innate immune cell populations. Changes in T-cells included increased CD8+ T-cell infiltration and expression of activation markers. It is currently unclear whether tumor-specific macrophages, dendritic cells, or extra-tumoral myeloid APCs are the primary drivers of CD8+ T-cell activation. It was apparent that the observed survival benefit did not depend on CD8+ T-cell changes, as neither selective antibody-mediated CD8+ T-cell depletion nor genetic T-cell diminished the survival benefit with treatment. It is unclear why increased CD8+ T-cell recruitment did not translate to an effect on survival in this tumor model. One possible explanation may lie in the fact that changes in the intratumor CD8+ T-cell population were not matched by the CD4+ T-cell population. Though CD8+ T-cells are widely viewed as the main effectors of anti-tumor adaptive immunity, CD4+ T-cells play pivotal roles. CD4+ T-cells are a diverse population with both anti-tumor and pro-tumor functions.^31,32^ For instance, Th1 CD4+ T-cells promote anti-tumor immunity in part by secreting IFNγ, which in turn facilitates CD8+ T-cell priming as well as antigen presentation.^31^ Though we did see a decrease in Treg infiltration, this alone may be insufficient to alter the immunosuppressive tumor microenvironment.^31^ Therefore, the apparent lack of CD8+ T-cell function may be explained by the lack of synchronous CD4+ T-cell infiltration or polarization toward immunogenic phenotypes. Alternatively, upregulation of other checkpoints such as TIGIT may be dominant in our model, and we are exploring this idea in ongoing experiments. Many other inhibitory signals, CXCL12/CXR4 signaling as one example, have been shown to dampen CD8+ T cell responses in the PDAC tumor microenvironment and we will seek to explore such possibilities in the future. Overall, these results show that increasing CD8+ T-cell infiltration alone is not sufficient to control tumor progression in these syngeneic orthotopic models of PDAC.

This study shows that the efficacy of combination SUMO inhibition and CD40 agonism in these orthotopic PDAC mouse models depends on macrophages. This is consistent with a prior report that showed CD40 agonism combined with gemcitabine also functions in a macrophage-dependent mechanism.^18^ Macrophage activation by anti-CD40 has been previously reported to require the presence of endogenous IFNγ. Buhtoiarov et. al demonstrated that macrophage priming by anti-CD40 was significantly decreased in IFNγ knockout mice, and that the macrophages themselves were the main source of IFNγ, creating a positive feedback loop.^33^ Enhanced tumoricidal activity of anti-CD40 activated alveolar macrophages has also been reported in a murine lung cancer model and is also dependent on IFNγ to potentiate macrophage production of nitric oxide and Interleukin-12.^34^ In PDAC, IFNγ has been shown to be necessary for infiltrating monocyte/macrophage matrix metalloproteinase mediated stromal degradation.^35^ SUMO inhibition has a interferon-augmenting effect that causes pro-immunogenic changes in macrophages in a type I IFN dependent mechanism in non-PDAC pre-clinical models.^6^ Here, we also observed increases in interferon signaling in PDAC tumors treated with TAK-981, an effect further amplified when combined with CD40 agonism. These results therefore also show that inhibiting SUMO may be a valid strategy for stimulating tumors-specific macrophages in PDAC to respond to additional macrophage-dependent immunotherapy. Because MHC-I downregulation is a well-established means of immune evasion,^9–13^ we were also interested in finding whether SUMO inhibition’s effects on interferon signaling coincided with increased MHC-I expression both *in vitro* and *in vivo.* In our case, TAK-981 primed tumors to respond to agonistic αCD40 to upregulate cancer specific MHC-I expression, even though basal levels of MHC-I expression both *in vitro* and *in vivo* are nearly undetectable in KPC46. Overall, TAK-981 appears to increase interferon signaling in macrophages, which in turn sensitizes them to further respond to CD40 agonism, and these combinatorial effects may play important roles in cancer-specific MHC-I expression and CD8+ T-cell infiltration and activation.

We hypothesize that TAK-981’s positive affect on interferon signaling by macrophages is critical for treatment efficacy in our tumor model, though this is subject to further study. In addition to the upregulation of IFNα and IFNγ in macrophages caused by combination therapy, our bioinformatic analysis showed that an additional possible mechanism contributing to increased macrophage activation is via decreased TGFβ signaling among macrophages, cancer cells, and fibroblasts. The upregulation of IFNγ signaling in macrophages could be linked to decreased TGFβ signaling, considering that these two pathways can negatively regulate each other. IFNγ has been shown to inhibit TGFβ through SMAD7, and TGFβ has been shown to inhibit IFNγ through MEK/ERK.^36^

The present study has several limitations. First, much of our work was done using a highly aggressive orthotopic model with a short median survival and lower basal levels of T-cell infiltration, as compared to published data using a similar orthotopic model.^13,26^ Though utilizing a highly aggressive model may offer assurances that our pre-clinical data may translate in clinical trials, our results may nonetheless be skewed. Secondly, though we have identified important changes in the PDAC TIME caused by SUMO inhibition plus CD40 agonism, and that this is driven by a macrophage-dependent mechanism, we have yet to decipher how this mechanistically relates to changes in T-cell infiltration. It is also puzzling that the influx of CD8+ T-cells does not appear to coincide with an increase in their activation, nor have we identified a strategy for doing this. Gaining a better insight of the complex interactions between innate and adaptive immunity in these tumor models may further guide us to develop new rationale combinations to better control PDAC growth in the laboratory and ultimately in the clinic.

## Conclusions

Inhibiting SUMOylation via the small molecule TAK-981, in combination with CD40 agonism, fundamentally alters the PDAC TIME by reprograming macrophages and inducing CD8+ T-cell infiltration. TAK-981 potentiates increases in interferon signaling in macrophages and T-cells, and CD40 provides a further additive affect. These in turn lead to a macrophage-dependent improvement in overall survival in a highly aggressive orthotopic mouse model of PDAC. These findings highlight a novel method to modulate a characteristically, highly immunosuppressive TIME. SUMO inhibition plus CD40 agonism warrants further investigation as a therapeutic combination in pancreatic cancer.

## Acknowledgements

We received support from the following organizations, grants, and gifts: Society for University Surgeons Resident Research Award (KYL) and Research for a Cure of Pancreatic Cancer (AML). TAK-981 used in the study was provided by Takeda Development Center Americas, Inc. We also received generous assistance from the following facilities: University of California San Diego (UCSD) Histopathology Core, UCSD Microscopy Core, UCSD IGM Genomics Center, La Jolla Institute for Immunology (LJII) Histopathology Core, and LJI Flow Cytometry Core.

## Figure Legends

**Supplementary Figure 1.**
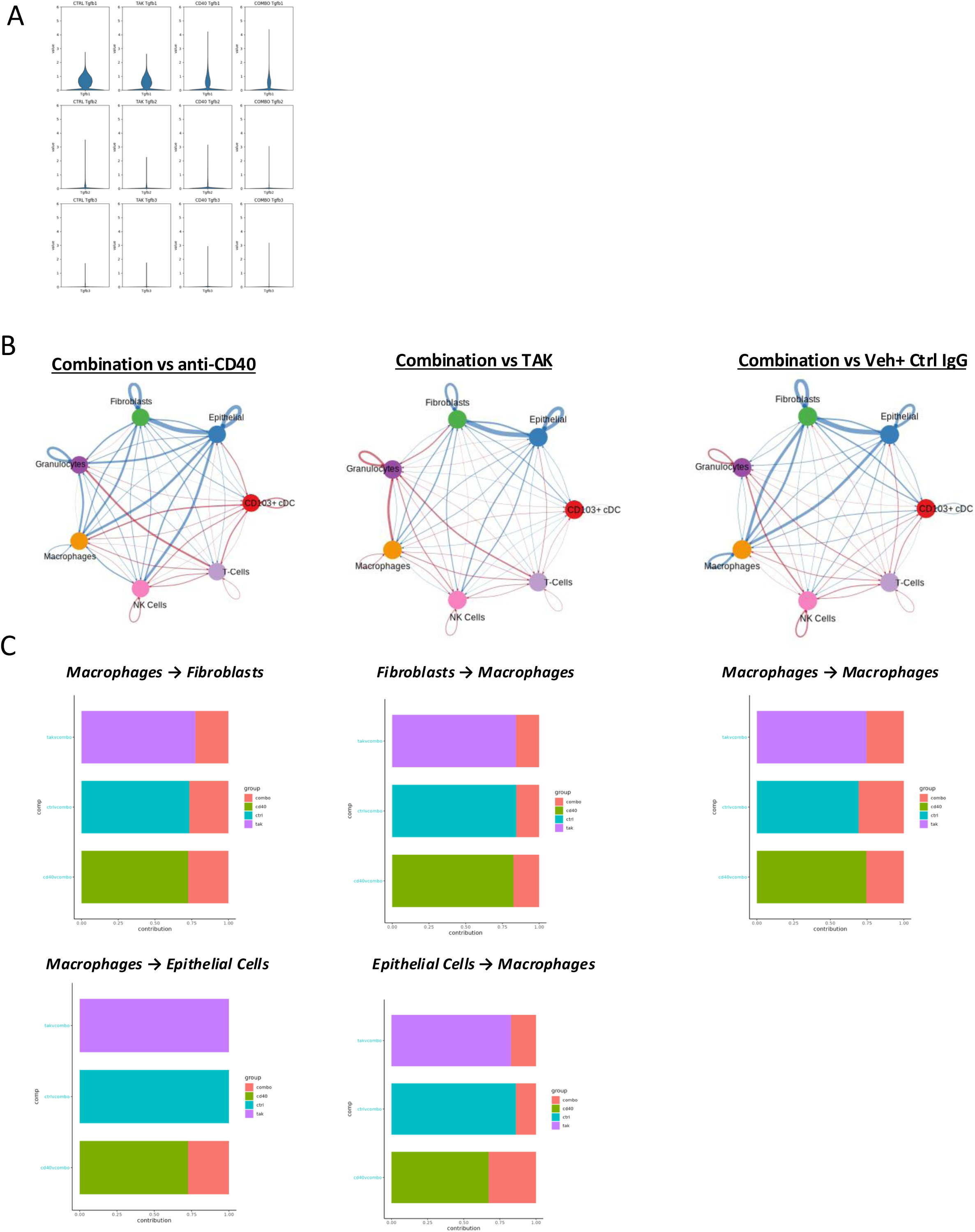
TAK-981 + anti-CD40 decreases TGFβ Signaling. A) Violin plots for TGBβ1 (combination vs αCD40: −0.35, combination vs TAK-981: −0.72, combination vs vehicle + IgG: −0.93, p<0.001 for all pairwise comparisons) and TGFβ2(combination vs anti-CD40: −1.35, combination vs TAK-981: −0.76, combination vs vehicle + IgG: −1.22, p<0.001 for all pairwise comparisons) differential expression coefficient levels in the combination treatment compared to monotherapy and untreated groups. B) Cell chat graphs showing decreased macrophage/epithelial and macrophage fibroblast interactions for combination vs αCD40, combination vs TAK-981, and combination vs vehicle + IgG conditions. C) Relative TGFβ information flow between epithelial cells/macrophages, fibroblasts/macrophages, and macrophages/macrophages for each of the following pairwise comparisons: combination vs αCD40, combination vs TAK-981, combination vs vehicle + IgG. For the following conditions, there were significant decreases in TGFβ signaling for all pairwise comparisons: macrophages vs. macrophages, macrophages vs. fibroblasts, fibroblasts vs. macrophages, and epithelial cells vs. macrophages.

**Supplemental Figure 2.**
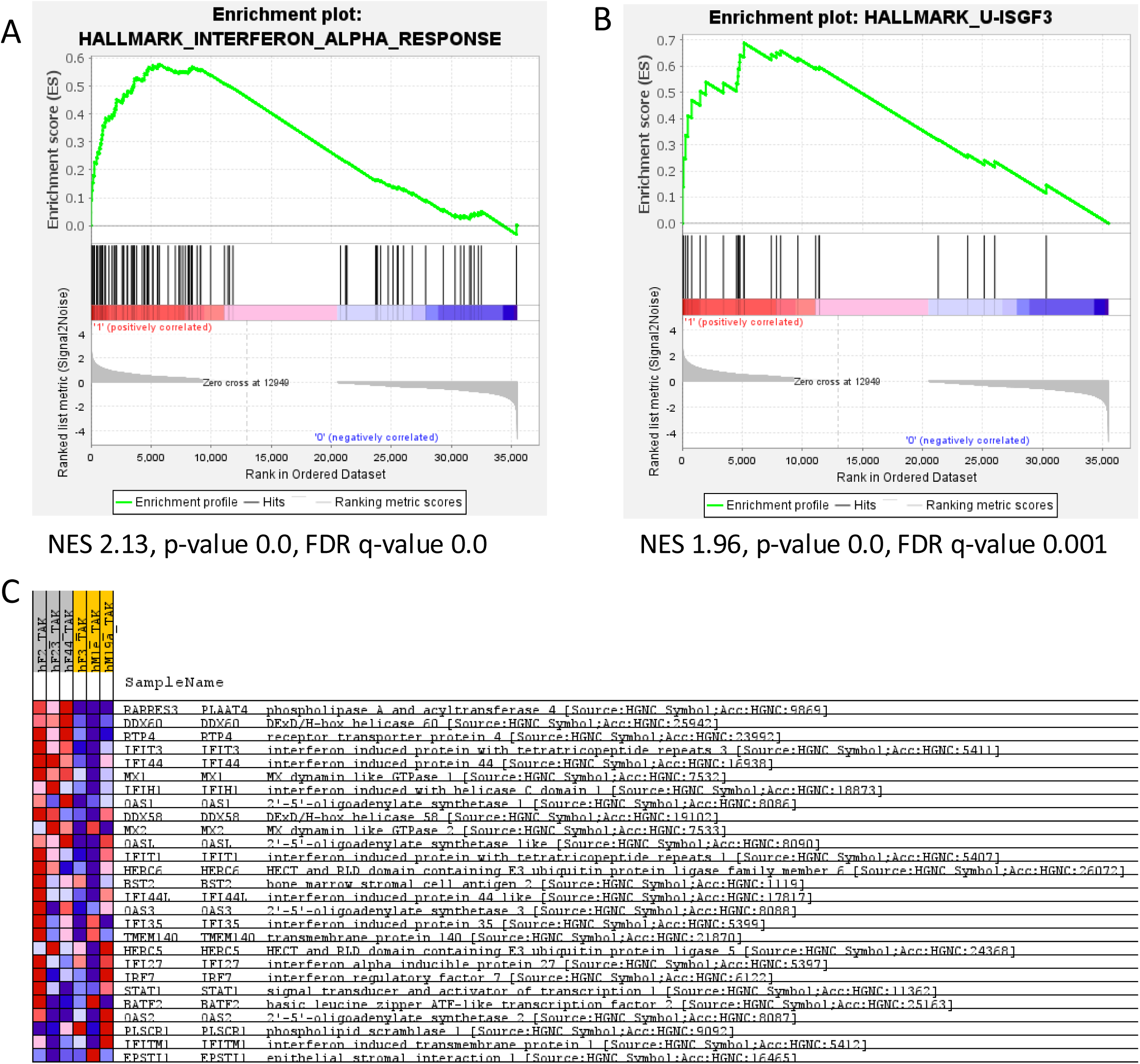
Sensitivity toward TAK-981 in patient derived organoids correlates with type I interferon pathway activation. A) GSEA leading edge plot comparing treated PDOs based on sensitivity to TAK-981 *in vitro* treatment. Utilized was the Hallmark Interferon Alpha Response gene set. B) GSEA leading edge plot utilizing genes previously classified as responsive to unphosphorylated ISGF3. Treated PDOs were divided based on sensitivity to TAK-981 *in vitro* treatment and compared. C) Heatmap corresponding to GSEA shown in panel B.

**Supplementary Figure 3.**
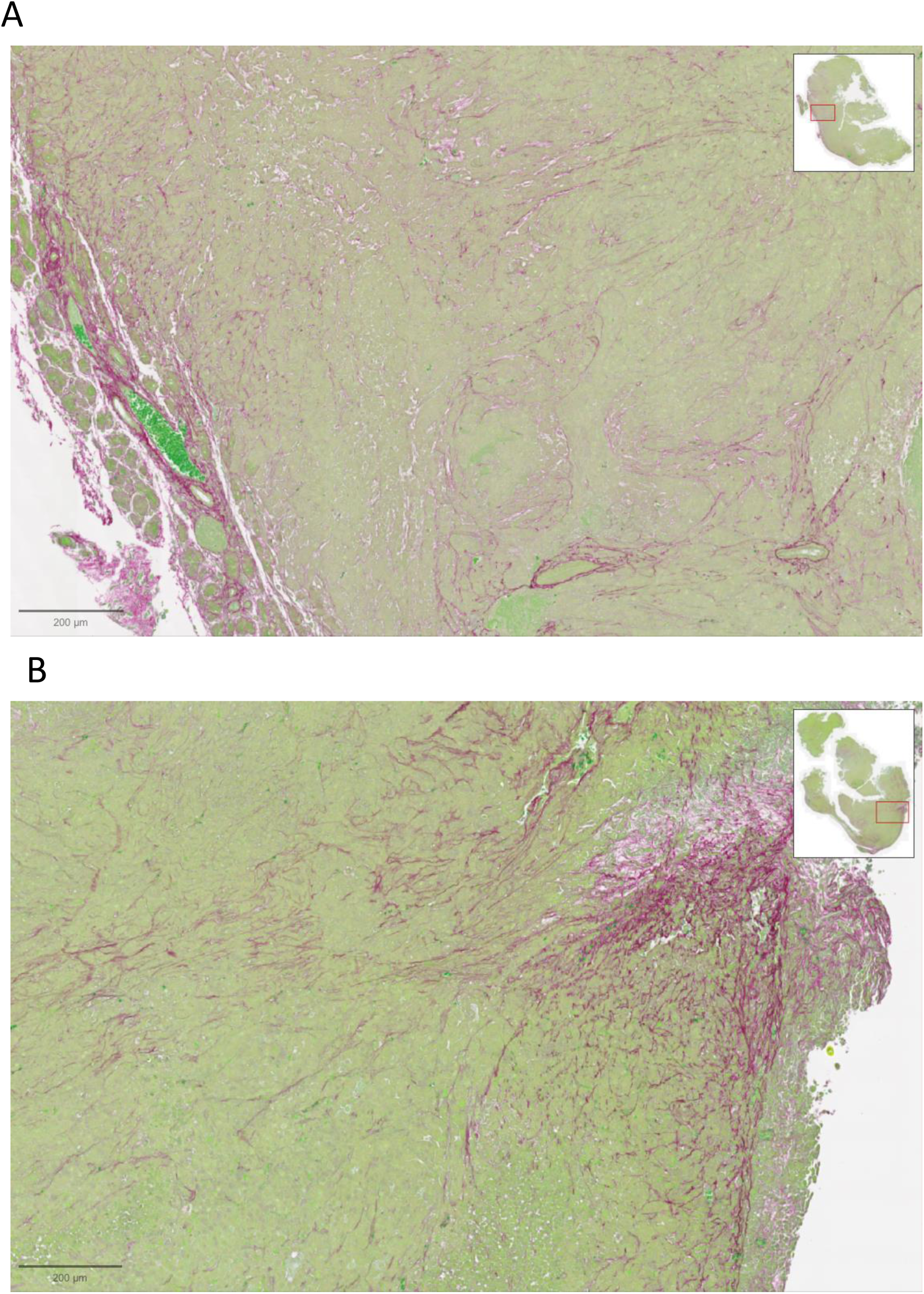
TAK-981 + aCD40 combination therapy does not affect collagen deposition as assessed by histologic staining. Tumors either treated with vehicle and isotype control (A) or combination TAK-981 + aCD40 (B) were stained for collagen with Sirius Red/Fast Green staining. There were no qualitative or quantitative differences in the amount of collagen staining.

**Supplementary Figure 4.**
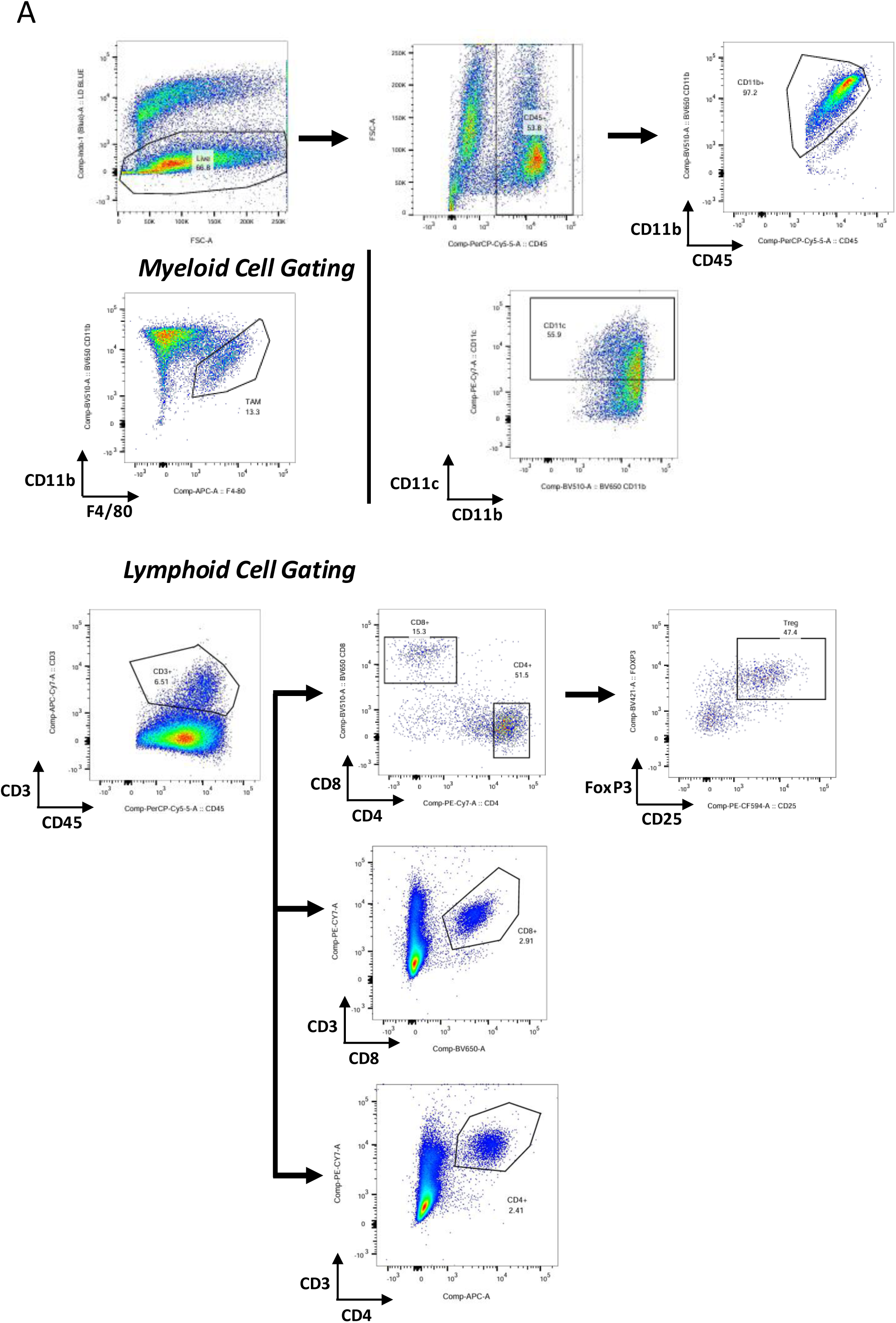
Flow Cytometry Gating Strategies. Example plots of gating schemes used in FlowJo to gate the following immune populations: CD8 T Cells(%CD3+/CD8+), CD4 T Cells(%CD3+/CD4+), Tregs (%CD4+/CD25+/FoxP3+), Macrophages(%CD11b+/F4/80+), Resident Macrophages(%CD11b+/F4/80+/CD103+), Dendritic Cells(%CD11b/CD11c+)

**Supplementary Figure 5.**
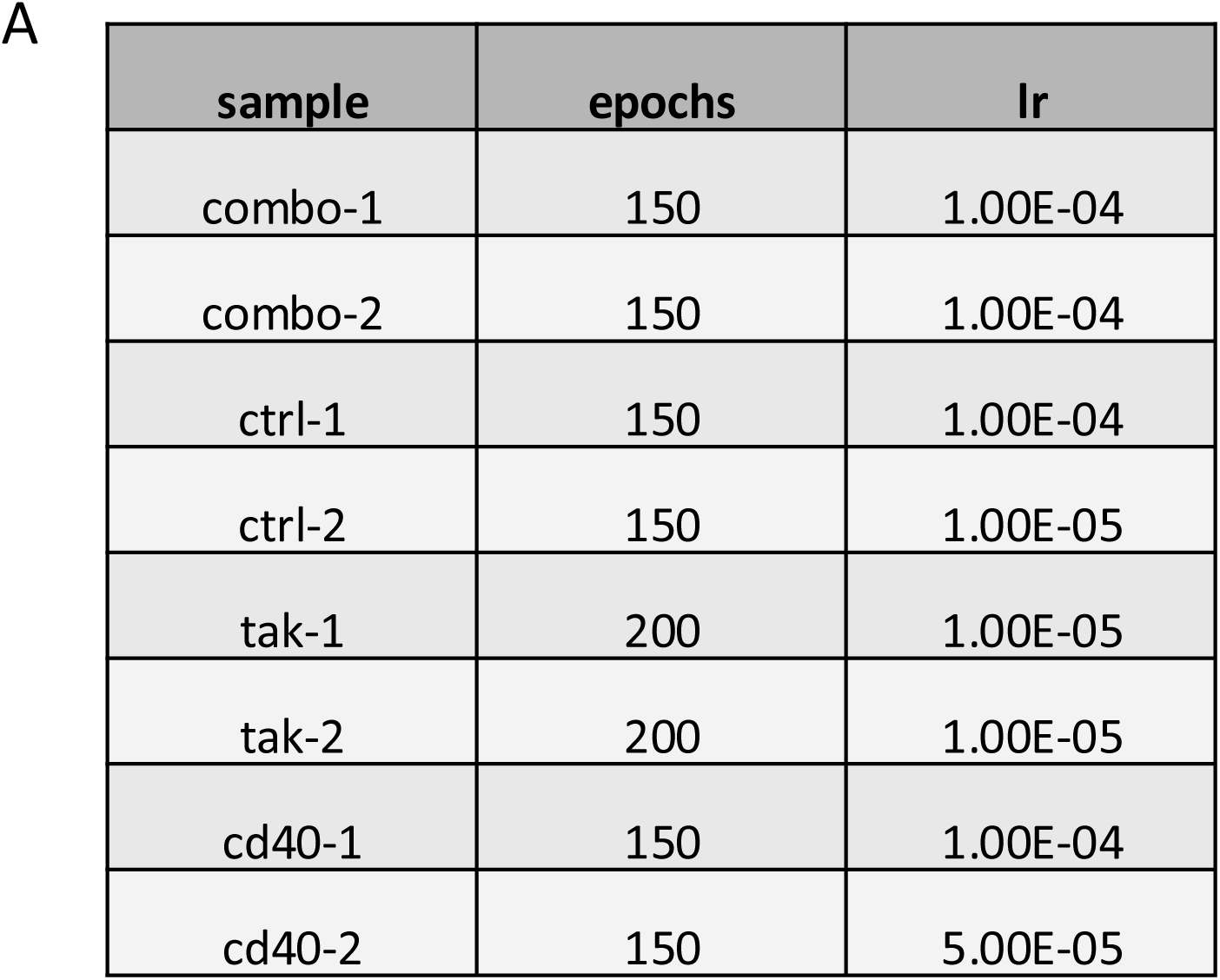
Per Sample Parameters for Cell Bender Training.

## Materials and Methods

### Cell lines and culture

The mouse PDAC cell line KPC46 was derived from a B6.129SF/J male *KPC* mouse. KPC cells were maintained in Roswell Park Memorial Institute (RPMI) 1640 (Sigmal-Aldrich, R8758) with 10% fetal bovine serum (FBS) and penicillin/streptomycin at 37°C with 5% CO_2_. The human cell line 79E was derived from a patient as previously described and maintained in RPMI-1640 with 10% FBS and penicillin/streptomycin at 37°C with 5% CO_2_[1].

### Organoid culture

Patient-derived organoids were generously given by the Tuveson Laboratory and maintained in a complete human medium, propagated, and sustained as previously described [2].

### PCR

Organoids were either grown in TAK-981 free media or pre-treated in 100nM of TAK-981 for 48hours and subsequently exposed to IFNα for either 3h, 6h, or 9h. KPC 46 or 79E cells were incubated in 1µM TAK-981 for either 2h, 4h, or 6h. RNA was then extracted using the RNeasy Plus Universal Mini it (Quiagen, Cat No:73404) and cDNA was generated using the iScript cDNA Synthesis Kit (BioRad, Cat. No: 1708890). PCR for STAT1, Mx1, Mx2, and Ifit2 was conducted using primers from IDT Technologies and the SSO Fast Evagreen PCR Kit (Biorad, Cat. No: 1725200). Statistical analysis was conducted using an unpaired two-tailed t test.

### Immunohistochemistry/Immunofluoresence

Tumor specimens were fixed in 10% NBF for 24 hours and then transferred to a 70% ethanol solution for an additional 24 hours. Tissue was then embedded in parrafin and sectioned. ***Immunohistochemistry:*** Slides were baked overnight at 65°C and subsequently cleared and re-hydrated. Antigen retrieval was performed using IHC Antigen Retrieval Solution-High pH(Invitrogen, Cat No:00-4956-56). Slides were then blocked in Bloxall (Vector, Cat. No: SP-6000) for 15 min and then a solution of 10% donkey/goat serum and 5% BSA for 1 hour. Slides were incubated overnight at 4°C in primary antibody: F4/80(1:150, Cell Signaling, Cat. No.:70076), slides were incubated in secondary antibody for 1hr at room temperature: goat anti-Rabbit (ImmPRESS HRP Goat Anti-Rabbit IgG Polymer Detection Kit, Vector Laboratories, Cat. No.: MP-7451). Immunohistochemistry slides were developed using ImmPact DAB Brown (Vector Labs, Cat. No.: SK-4105). ***Immunofluoresence:*** Slides were baked at 60°C for 1 hour and then cleared and re-hydrated. Staining was conducted using the Intellipath Automated IHC Stainer (Biocare medical). Antigen retrieval was performed using Antigen Unmasking Solution (Citrate Based, pH6) (Vector, H-3300). Slides were then blocked using Bloxall(Vector, Cat. No: SP-6000) for 10 min and Blotto solution (Thermo, PI37530) for 10 min.

Staining for CD8α and CD4 was conducted using the following antibodies: CD8α - Anti-CD8 (Rat, Invitrogen, 14-0195-82, 1:50) for 1 h, Anti-Rat HRP Polymer (Cell IDX, 2AH-50) for 30 min, Opal 570 (Akoya, FP1488001KT) for 10 min; CD4- Anti-CD4 (Rabbit, Abcam, ab288724, 1:1000) for 1 hr, Anti-Rabbit HRP Polymer (Cell IDX, 2RH-50) for 30 min, Opal 690 (Akoya, FP1497001KT) for 10 min. Nuclei were stained using DAPI (1µg/ml) for 15 min.

### Flow Cytometry

#### KPC 46 and KPC1199 Flow Cytometry for MHCI

Cells were plated and maintained until approximately 50-75% confluence in Roswell Park Memorial Institute (RPMI) 1640 (Sigmal-Aldrich, R8758) with 10% fetal bovine serum (FBS) and penicillin/streptomycin at 37°C with 5% CO_2_. At that time, they were dosed with 100 nM TAK-981 or left untreated. Approximately 24 hours later, cell cultures selected to receive IFNg were exposed to 100 U/mL of IFNg for 24 hours. After this incubation, cells were harvested for flow cytometry. Cells were washed with cold PBS, labeled with viability dye (Zombie Yellow, BioLegend), and labeled with anti-MHC-I fluorescent antibody (anti-mouse MHCH-2kd/H-2Dd, BioLegend, Clone: 34-1-25). Cells were then washed with PBS and analyzed with the NovoCyte Advanteon Flow Cytometer (Agilent, Santa Clara, CA) according to manufacturer protocols.

#### Dissociation of Tumor and Spleen Tissue and Flow Cytometry

Spleens were collected and processed into single-cell suspension by mashing the spleen through a 70-μm filter with the back of a syringe and washing the filter with 10 mL of PBS 2% FBS. Splenocytes were spun at 300 × *g* for 10 minutes and then subjected to Red Blood Cell lysis (RBC) using 5 ml of BD Pharmalyse (BD). After 5 minutes of incubation, splenocytes were washed with 5 ml of PBS 2%FBS and counted. Tumors were minced into small pieces and incubated at 37°C with rotation for 30 minutes in 5 mL of digestion buffer consisting on DMEM high glucose, 10% Gentle Collagenase/Hyaluronidase (GCH, STEM CELL), 10% FBS, and 10% DNaseI (1 mg/mL stock, Roche). Samples were then added on top of a 70-μm filter and smashed with the back of a syringe while adding 5 mL of PBS 2% FBS. Flow through was spun at 300 × *g* for 10 minutes and subjected to RBC lysis as described above using 3 ml of RBC lysis buffer. Cells were counted and subjected to magnetic separation of tumor infiltrating lymphocytes using Easysep TIL kit (STEM CELL). The flow through was collected for epithelial cell analysis of MHC-I while the purified cells were stained for immunophenotyping. After 15 minutes of incubation with FcBlock in 50ul, extracellular antibody mix in brilliant buffer (BD) was added and incubated for 30 min. After washing, LD blue fixable viability dye (Invitrogen) was added and incubated for 30 min. Samples were then washed, fixed and permeabilized with FoxP3 transcription factor staining buffer (Thermo Fisher) and intracellular antibodies were added and incubated overnight at 4°C. Samples were run on an LSR fortessa and analyzed with FlowJo^TM^ (Becton, Dickinson & Company).

Extracellular antibodies for flow cytometry: Anti-CD45 (Biolegend, Clone: 30-F11), Anti-CD44 (Biolegend, Clone: IM7), Anti-CD62L (BD Bioscences, Clone: MEL-14), Anti-CD4 (Biolegend, Clone: GK1.5), Anti-PD1 (Biolegend, Clone: 29F.1A12), Anti-CD8 (Biolegend, Clone: 53-6.7), Anti-CTLA-4 (Biolegend, Clone: UC10-4B9), Anti-CD3 (Biolegend, Clone: 17A2), Anti-CD25 (Biolegend, Clone: PC61), Anti-CD103 (Biolegend, Clone: 2E7), Anti-CD69 (BD Biosciences, Clone: H1.2F3), Anti-F4/80 (Biolegend, Clone:BM8), Anti-CD11b (Biolegend, Clone: M1/70), Anti-CD11c (Biolegend, Clone: HL3), Anti-MHCII (Biolegend, Clone: M5/114.15.2), Anti-EPCAM(Biolegend, Clone: G8.8).

Intracellular antibodies for flow cytometry: Anti-FoxP3 (Biolegend, Clone: MF-14), Anti-CD206 (Biolegend, Clone:C068C2).

### Animal models and in vivo model generation

Animals were raised and experimented on according to protocols approved by the University of California San Diego Institutional Animal Care and Use Committee (protocol number S09158). B6.129SF/J and B6 Nude mice were purchased from The Jackson Laboratory.

### Animal survival experiments

PDAC tumors were orthotopically generated as follows: 2,000 KPC46 cells resuspended in 20 µL Matrigel (Corning Matrigel Matrix, Cat. No.: 356231) were directly injected into the pancreatic tail of mice. The mice were anesthetized with 100 mg/kg ketamine and 10 mg/kg xylazine. The mice were monitored for 5 days post-operatively. Tumor burden was established by serial ultrasonography (Convex L20 HD3, Clarius). Mice were randomized into treatment groups after tumors reached at least 3-5 mm in diameter as measured by ultrasound. TAK-981 in lyophilized powder form was resuspended water with 5% dextrose and 20% Kolliphor (Sigma-Aldrich, Cat. No: 30906). Mice were dosed with 15 mg/kg TAK-981 (Takeda Pharmaceuticals) every 48 hours via IP injection following enrollment. Anti-CD40 antibody was obtained from BioXCell (BioXCell clone FGK4.5/FGK45, Cat. No.: BP0016-2) and diluted in a pH 7.0 dilution buffer supplied by BioXCell (Cat. No. IP0070) according to manufacturer instructions. Mice were dosed with 100µg anti-CD40 antibody weekly via IP injection. Mice treated with both drugs were first pre-treated with TAK-981 for one dose, and then two days later were administered a subsequent dose of TAK-981 and a dose of anti-CD40 antibody. Treatments were continued on this schedule until the time of death or moribund status prompting euthanasia, according to IACUC protocols. Statistical analysis comparing survival was conducted using a log rank test.

### CD8 T Cell Depletion

Orthotopic tumors were established in B6.129SF1/J mice (strain 101043; Jackson Laboratory, Ben Harbor, Maine) by injecting 2,000 individual KPC46 cells into the distal pancreas of these mice, as described previously. Tumors were allowed to grow until they were detectable on ultrasound. Mice were randomized into groups receiving anti-CD8 antibody (anti-mouse CD8 antibody, clone YTS 169.4, BioXCell, Lebanon, NH) and those receiving KLH isotype controls (anti-KLH, clone LTF-2, Lebanon, NH). Both antibodies were administered every other day at a dose of 200 µg per mouse, inject intraperitoneally. These injections occurred regularly up until the mouse’s euthanasia or death.

### Liposomal Clodronate Macrophage Depletion

PDAC tumor were generated as previously described above and mice were enrolled when tumor diameter reached 3-5mm, as assessed by ultrasound. Mice enrolled in TAK-981 and anti-CD40 combination treatment groups were dosed with either 200 µl clodronate liposome (Clodrosome, Encapsula Nanosciences, Cat No: CLD-8909) or PBS liposome (Encapsome, Encapsula Nanosciences, Cat. No: CLD-8910) via IP injection every four days. Two days after macrophage depletion was initiated, TAK-981 and anti-CD40 treatment administration began, as described above. Both Clodronate liposome and PBS liposome administration were continued on this schedule until the time of death or moribund status prompting euthanasia, according to IACUC protocols.

### Bulk RNA Sequencing

For bulk RNA-seq experiments, organoids treated with 100 nM TAK981 for 48 hrs in Matrigel were lysed directly with 1 mL of TRIzol reagent (Thermo Fisher), and total RNA was extracted according to the manufacturer’s instructions. RNA-seq libraries were constructed using according to the manufacturer’s instructions (Illumina). Briefly, 2 μg of purified RNA was poly-A selected and fragmented with fragmentation enzyme. cDNA was synthesized with Super Script II master mix, followed by end repair, A-tailing, and PCR amplification. RNA-seq libraries were sequenced using an Illumina platform.

The organoids included in the bulk RNA sequencing experiment were hF2, hF3, hF23, hF44, which are from pancreas, hM1e from lung, and hM19A from liver. We have also categorized hF2, hF23, and hF44 as the sensitive organoids that are sensitive to TAK-981 treatment and hF3, hM1e, hM19A as the resistant organoids. The data was processed and quantified using kallisto[3]. The processed data was analyzed with GSEA Software[4]. For the reference gene sets, Hallmark gene sets from MSigDB was used[5]. To calculate the normalized enrichment score (NES), signal2Noise metrics was used for categorical phenotypes. To filter out insignificant results, I set the minimum number of gene sets used for analysis as 10 and maximum as 500[6]. The number of permutation was 10,000 and the maximum expression value for the identifiers was used.

### Single Cell RNA Sequencing

For single-cell RNA library preparation, 2 mouse tumor samples per treatment group were randomly selected and 20,000 cells from each tumor sample were isolated. Cell barcoding, RT, cDNA amplification and library construction were done using Chromium Next GEM Single Cell 3ʹ HT v3.1 (10x Genomics). The libraries were sequenced on Illumina NovaSeq 6000 with the S4 kit at 1 billion reads per sample.

### QC and Preprocessing

Data was processed using Cellranger 8.0.1 and aligned to GRCm39-2024-A using default settings. Raw matrices were inputted into cellbender(v0.3.0) and processed using default settings[7]. If training failed to converge, epochs or learning rate was adjusted. Per sample training parameters are in (SUPP TABLE). Droplets with a probability of 0.5 or greater were selected. Doublets were then removed using vaeda (v0.0.30)[8]. Cells were then filtered in scanpy (v1.10.1), removing cells with high mitochondrial content, and low counts[9].

### Clustering

Cells were then integrated using scvi-tools (v1.2.0)[10]. Briefly, the top 8000 most variable genes were selected using Poisson gene selection in scvi. Then, correcting for sample effects as well as ribosomal content, an autotune run was conducted to select the best parameters. Using these parameters, the model was then trained for 58 epochs. The neighborhood space was computed off of this, using 30 neighbors. UMAP was then computed using a minimum distance of 0.1. Cells were then clustered using leiden in scanpy, using the igraph flavor, 2 iterations, and a resolution of 0.4. Marker genes were computed using the Wilcoxon rank sum test, and cells were annotated based on marker gene expression.

### Differential Gene expression

Differential gene expression was done using memento (v0.1.0), using a variable for samples as well as treatment, and bootstrapped for 10000 iterations[11].

### Differential Abundance

The scCODA implementation in pertpy (v0.9.4) was used to test for differences in cell composition[12,13]. All tests were done using Epithelial cells as the reference population.

### Gene Set Enrichment Analysis

GSEA was done using clusterprofiler against the Hallmark gene sets in msigDB[14].

### CellChat

CellChat was done as described in the tool authors publication[15]. Only secreted signaling was analyzed

## References

1. Conroy T, Hammel P, Hebbar M, et al. FOLFIRINOX or Gemcitabine as Adjuvant Therapy for Pancreatic Cancer. N Engl J Med. 2018;379(25):2395–2406. doi:10.1056/NEJMoa1809775

2. Neoptolemos JP, Palmer DH, Ghaneh P, et al. Comparison of adjuvant gemcitabine and capecitabine with gemcitabine monotherapy in patients with resected pancreatic cancer (ESPAC-4): a multicentre, open-label, randomised, phase 3 trial. The Lancet. 2017;389(10073):1011–1024. doi:10.1016/S0140-6736(16)32409-6

3. Balachandran VP, Beatty GL, Dougan SK. Broadening the Impact of Immunotherapy to Pancreatic Cancer: Challenges and Opportunities. Gastroenterology. 2019;156(7):2056–2072. doi:10.1053/j.gastro.2018.12.038

4. Decque A, Joffre O, Magalhaes JG, et al. Sumoylation coordinates the repression of inflammatory and anti-viral gene-expression programs during innate sensing. Nat Immunol. 2016;17(2):140–149. doi:10.1038/ni.3342

5. Lightcap ES, Yu P, Grossman S, et al. A small-molecule SUMOylation inhibitor activates antitumor immune responses and potentiates immune therapies in preclinical models. Sci Transl Med. 2021;13(611):eaba7791. doi:10.1126/scitranslmed.aba7791

6. Nakamura A, Grossman S, Song K, et al. The SUMOylation inhibitor subasumstat potentiates rituximab activity by IFN1-dependent macrophage and NK cell stimulation. Blood. 2022;139(18):2770–2781. doi:10.1182/blood.2021014267

7. Kumar S, Schoonderwoerd MJA, Kroonen JS, et al. Targeting pancreatic cancer by TAK-981: a SUMOylation inhibitor that activates the immune system and blocks cancer cell cycle progression in a preclinical model. Gut. Published online January 24, 2022:gutjnl-2021-324834. doi:10.1136/gutjnl-2021-324834

8. Erdem S, Lee HJ, Shankara Narayanan JSN, et al. Inhibition of SUMOylation Induces Adaptive Antitumor Immunity against Pancreatic Cancer through Multiple Effects on the Tumor Microenvironment. Mol Cancer Ther. 2024;23(11):1597–1612. doi:10.1158/1535-7163.MCT-23-0572

9. Demel UM, Böger M, Yousefian S, et al. Activated SUMOylation restricts MHC class I antigen presentation to confer immune evasion in cancer. Journal of Clinical Investigation. 2022;132(9):e152383. doi:10.1172/JCI152383

10. Pandha H, Rigg A, John J, Lemoine N. Loss of expression of antigen-presenting molecules in human pancreatic cancer and pancreatic cancer cell lines. Clinical and Experimental Immunology. 2007;148(1):127–135. doi:10.1111/j.1365-2249.2006.03289.x

11. Ryschich E, Nötzel T, Hinz U, et al. Control of T-cell-mediated immune response by HLA class I in human pancreatic carcinoma. Clin Cancer Res. 2005;11(2 Pt 1):498–504.

12. Sharma P, Hu-Lieskovan S, Wargo JA, Ribas A. Primary, Adaptive, and Acquired Resistance to Cancer Immunotherapy. Cell. 2017;168(4):707–723. doi:10.1016/j.cell.2017.01.017

13. Yamamoto K, Venida A, Yano J, et al. Autophagy promotes immune evasion of pancreatic cancer by degrading MHC-I. Nature. 2020;581(7806):100–105. doi:10.1038/s41586-020-2229-5

14. Vonderheide RH. CD40 Agonist Antibodies in Cancer Immunotherapy. Annu Rev Med. 2020;71(1):47–58. doi:10.1146/annurev-med-062518-045435

15. Vonderheide RH. The Immune Revolution: A Case for Priming, Not Checkpoint. Cancer Cell. 2018;33(4):563–569. doi:10.1016/j.ccell.2018.03.008

16. Khalil M, Vonderheide RH. Anti-CD40 agonist antibodies: Preclinical and clinical experience. Update on Cancer Therapeutics. 2007;2(2):61–65. doi:10.1016/j.uct.2007.06.001

17. Salomon R, Dahan R. Next Generation CD40 Agonistic Antibodies for Cancer Immunotherapy. Front Immunol. 2022;13:940674. doi:10.3389/fimmu.2022.940674

18. Beatty GL, Chiorean EG, Fishman MP, et al. CD40 Agonists Alter Tumor Stroma and Show Efficacy Against Pancreatic Carcinoma in Mice and Humans. Science. 2011;331(6024):1612–1616. doi:10.1126/science.1198443

19. Byrne KT, Betts CB, Mick R, et al. Neoadjuvant Selicrelumab, an Agonist CD40 Antibody, Induces Changes in the Tumor Microenvironment in Patients with Resectable Pancreatic Cancer. Clin Cancer Res. 2021;27(16):4574–4586. doi:10.1158/1078-0432.CCR-21-1047

20. Long KB, Gladney WL, Tooker GM, Graham K, Fraietta JA, Beatty GL. IFNγ and CCL2 Cooperate to Redirect Tumor-Infiltrating Monocytes to Degrade Fibrosis and Enhance Chemotherapy Efficacy in Pancreatic Carcinoma. Cancer Discovery. 2016;6(4):400–413. doi:10.1158/2159-8290.CD-15-1032

21. Baker LA, Tiriac H, Clevers H, Tuveson DA. Modeling Pancreatic Cancer with Organoids. Trends in Cancer. 2016;2(4):176–190. doi:10.1016/j.trecan.2016.03.004

22. Tiriac H, Belleau P, Engle DD, et al. Organoid Profiling Identifies Common Responders to Chemotherapy in Pancreatic Cancer. Cancer Discov. 2018;8(9):1112–1129. doi:10.1158/2159-8290.CD-18-0349

23. Vitale G, van Eijck CHJ, van Koetsveld Ing PM, et al. Type I Interferons in the Treatment of Pancreatic Cancer: Mechanisms of Action and Role of Related Receptors. Annals of Surgery. 2007;246(2):259–268. doi:10.1097/01.sla.0000261460.07110.f2

24. Kotredes KP, Gamero AM. Interferons as Inducers of Apoptosis in Malignant Cells. J Interferon Cytokine Res. 2013;33(4):162–170. doi:10.1089/jir.2012.0110

25. Subramanian A, Tamayo P, Mootha VK, et al. Gene set enrichment analysis: A knowledge-based approach for interpreting genome-wide expression profiles. Proceedings of the National Academy of Sciences. 2005;102(43):15545–15550. doi:10.1073/pnas.0506580102

26. Datta J, Dai X, Bianchi A, et al. Combined MEK and STAT3 Inhibition Uncovers Stromal Plasticity by Enriching for Cancer-Associated Fibroblasts With Mesenchymal Stem Cell-Like Features to Overcome Immunotherapy Resistance in Pancreatic Cancer. Gastroenterology. 2022;163(6):1593–1612. doi:10.1053/j.gastro.2022.07.076

27. Garg B, Giri B, Modi S, et al. NFκB in Pancreatic Stellate Cells Reduces Infiltration of Tumors by Cytotoxic T Cells and Killing of Cancer Cells, via Up-regulation of CXCL12. Gastroenterology. 2018;155(3):880–891.e8. doi:10.1053/j.gastro.2018.05.051

28. Ischenko I, D’Amico S, Rao M, et al. KRAS drives immune evasion in a genetic model of pancreatic cancer. Nat Commun. 2021;12(1):1482. doi:10.1038/s41467-021-21736-w

29. O’Reilly EM, Oh DY, Dhani N, et al. Durvalumab With or Without Tremelimumab for Patients With Metastatic Pancreatic Ductal Adenocarcinoma: A Phase 2 Randomized Clinical Trial. JAMA Oncology. 2019;5(10):1431–1438. doi:10.1001/jamaoncol.2019.1588

30. Royal RE, Levy C, Turner K, et al. Phase 2 Trial of Single Agent Ipilimumab (Anti-CTLA-4) for Locally Advanced or Metastatic Pancreatic Adenocarcinoma. Journal of Immunotherapy. 2010;33(8):828–833. doi:10.1097/CJI.0b013e3181eec14c

31. Kim HJ, Cantor H. CD4 T-cell Subsets and Tumor Immunity: The Helpful and the Not-so-Helpful. Cancer Immunology Research. 2014;2(2):91–98. doi:10.1158/2326-6066.CIR-13-0216

32. Borst J, Ahrends T, Bąbała N, Melief CJM, Kastenmüller W. CD4+ T cell help in cancer immunology and immunotherapy. Nat Rev Immunol. 2018;18(10):635–647. doi:10.1038/s41577-018-0044-0

33. Buhtoiarov IN, Lum H, Berke G, Paulnock DM, Sondel PM, Rakhmilevich AL. CD40 Ligation Activates Murine Macrophages via an IFN-γ-Dependent Mechanism Resulting in Tumor Cell Destruction In Vitro 1. The Journal of Immunology. 2005;174(10):6013–6022. doi:10.4049/jimmunol.174.10.6013

34. Imaizumi K, Kawabe T, Ichiyama S, et al. Enhancement of tumoricidal activity of alveolar macrophages via CD40-CD40 ligand interaction. American Journal of Physiology-Lung Cellular and Molecular Physiology. 1999;277(1):L49–L57. doi:10.1152/ajplung.1999.277.1.L49

35. Derynck R, Turley SJ, Akhurst RJ. TGF-β biology in cancer progression and tumor immunotherapy. Nat Rev Clin Oncol. 2021;18(1):9–34. doi:10.1038/s41571-020-0403-1

36. Gauthier T, Chen W. IFN-γ and TGF-β, Crucial Players in Immune Responses: A Tribute to Howard Young. J Interferon Cytokine Res. 2022;42(12):643–654. doi:10.1089/jir.2022.0132

## References

1. Buckel L, Savariar EN, Crisp JL, et al. Tumor Radiosensitization by Monomethyl Auristatin E: Mechanism of Action and Targeted Delivery. Cancer Research. 2015;75(7):1376–1387. 10.1158/0008-5472.can-14-1931

2. Boj Sylvia F, Hwang CI, Baker Lindsey A, et al. Organoid Models of Human and Mouse Ductal Pancreatic Cancer. Cell. 2015;160(1-2):324–338. 10.1016/j.cell.2014.12.021

3. Bray NL, Pimentel H, Melsted P, Pachter L. Near-optimal probabilistic RNA-seq quantification. Nature Biotechnology. 2016;34(5):525–527. 10.1038/nbt.3519

4. Mootha VK, Lindgren CM, Eriksson KF, et al. PGC-1α-responsive genes involved in oxidative phosphorylation are coordinately downregulated in human diabetes. Nature Genetics. 2003;34(3):267–273. 10.1038/ng1180

5. Arthur Liberzon, Aravind Subramanian, Reid Pinchback, Helga Thorvaldsdóttir, Pablo Tamayo, Jill P. Mesirov, Molecular signatures database (MSigDB) 3.0, *Bioinformatics*, Volume 27, Issue 12, June 2011, Pages 1739–1740, 10.1093/bioinformatics/btr260

6. Liberzon A, Birger C, Thorvaldsdóttir H, Ghandi M, Mesirov Jill P, Tamayo P. The Molecular Signatures Database Hallmark Gene Set Collection. Cell Systems. 2015;1(6):417–425. 10.1016/j.cels.2015.12.004

7. Fleming SJ, Chaffin MD, Arduini A, et al. Unsupervised removal of systematic background noise from droplet-based single-cell experiments using CellBender. Nature Methods. 2023;20(9):1323–1335. 10.1038/s41592-023-01943-7

8. Schriever H. Vaeda computationally annotates doublets in single-cell RNA sequencing data | Bioinformatics | Oxford Academic. Accessed November 22, 2024. https://academic.oup.com/bioinformatics/article/39/1/btac720/6808614

9. Wolf FA, Angerer P, Theis FJ. SCANPY: large-scale single-cell gene expression data analysis. Genome Biology. 2018;19(1). 10.1186/s13059-017-1382-0

10. Gayoso A, Lopez R, Xing G, et al. A Python library for probabilistic analysis of single-cell omics data. Nature Biotechnology. 2022;40(2):163–166. 10.1038/s41587-021-01206-w

11. Kim MC, Gate R, Lee DS, et al. Method of moments framework for differential expression analysis of single-cell RNA sequencing data. Cell. 2024;187(22):6393–6410.e16. 10.1016/j.cell.2024.09.044

12. Büttner M, Johannes Ostner, Müller CL, Theis FJ, Schubert B. scCODA is a Bayesian model for compositional single-cell data analysis. Nature Communications. 2021;12(1). 10.1038/s41467-021-27150-6

13. Heumos L, Ji Y, May L, et al. Pertpy: an end-to-end framework for perturbation analysis. Published online August 7, 2024. 10.1101/2024.08.04.606516

14. Xu S, Hu E, Cai Y, et al. Using clusterProfiler to characterize multiomics data. Nature Protocols. Published online July 17, 2024. 10.1038/s41596-024-01020-z

15. Jin S, Plikus MV, Nie Q. CellChat for systematic analysis of cell–cell communication from single-cell transcriptomics. Nature Protocols. Published online September 16, 2024. 10.1038/s41596-024-01045-4

